# brisc: a multithreaded CPU toolkit for single-cell analysis

**DOI:** 10.64898/2026.11.03.764120

**Authors:** Keon Arbabi, Tain Luquez, Jaiz Jeeson, Mingyau Lam, Hamed Heydari, Gary D. Bader, Vilas Menon, Michael Wainberg

**Affiliations:** Lunenfeld-Tanenbaum Research Institute, Mount Sinai Hospital, Toronto, Canada; Institute of Medical Science, University of Toronto, Toronto, Canada; Center for Translational and Computational Neuroimmunology, Department of Neurology, Columbia University Irving Medical Center, New York, United States of America; The Donnelly Centre, University of Toronto, Toronto, Canada; Department of Molecular Genetics, University of Toronto, Toronto, Canada; Princess Margaret Research Institute, University Health Network, Toronto, Canada; Department of Computer Science, University of Toronto, Toronto, Canada; Department of Psychiatry, University of Toronto, Toronto, Canada; Division of Biostatistics, Dalla Lana School of Public Health, University of Toronto, Toronto, Canada

## Abstract

Single-cell RNA sequencing datasets have grown from tens of thousands of cells to tens of millions. At this scale, standard toolkits like Scanpy and Seurat need hours per analysis, hindering iterative exploration. We present brisc, the first end-to-end multithreaded CPU toolkit for single-cell analysis, spanning dimensionality reduction, integration, label transfer, clustering, embedding, differential expression and plotting. Unlike GPU-based counterparts, brisc strictly guarantees reproducibility: identical results between runs, regardless of thread count. Algorithmic innovations include a blocked inverted-file-index-based nearest-neighbors algorithm that makes 81–98% fewer errors than Scanpy and Seurat. On a 192-CPU node, brisc ran a standard workflow on 10 million cells in 3 minutes, 116× faster than Scanpy at under half its peak memory usage and 236× faster than Seurat with BPCells. On 96 CPUs, it matched the speed of rapids-singlecell on four H100 GPUs. brisc enables near-real-time, reproducible analysis of tens of millions of cells without specialized hardware.

## Introduction

Single-cell RNA sequencing has become a mainstay of biology, and its scale has grown exponentially. Experiments increasingly profile millions of cells, and atlas-scale efforts now exceed tens of millions. Scanpy^1^ and Seurat^2^, the field’s most popular toolkits, were designed when hundreds of thousands of cells was a large dataset, and remain largely single-threaded and memory-intensive. On today’s datasets, a single pass through the standard workflow (quality control, normalization, dimensionality reduction, clustering and embedding, among other steps) can take hours. Worse, analysis is rarely a single pass: quality-control thresholds are revisited, integration is tuned, and clustering (one of the slowest steps) is rerun at multiple resolutions. In practice, runtime constraints limit the scope of what analysts attempt.

A common practical solution is to analyze less data, either by sketching or by splitting. Sketching performs dimensionality reduction and clustering on a small, representative subset of cells, typically tens of thousands, then extrapolates the resulting clusters to the full dataset. Splitting divides the data, usually by batch or sample, and analyzes each portion separately. Because sketching and splitting identify cell types based on subsets of the data, they risk missing rare cell types that appear too infrequently in the subset to be detectable as a distinct cluster.

Two software engineering strategies have emerged to address scaling challenges directly (Fig. 1a). Out-of-core computation, used by toolkits like BPCells^3^ and Scarf^4^, reduces peak memory usage by keeping the count matrix on disk and streaming it through memory in small chunks. Seurat relies on BPCells as a backend for large datasets. Streaming from disk has two main limitations: inflexibility, since any operation not explicitly implemented by the toolkit requires loading the full dataset into memory first (which defeats the purpose of out-of-core computation); and runtime, since every step requires reloading the dataset from disk (sometimes dozens of times per step), which is several orders of magnitude slower than reading from memory, especially on shared computing clusters with networked file systems.

**Fig. 1.**
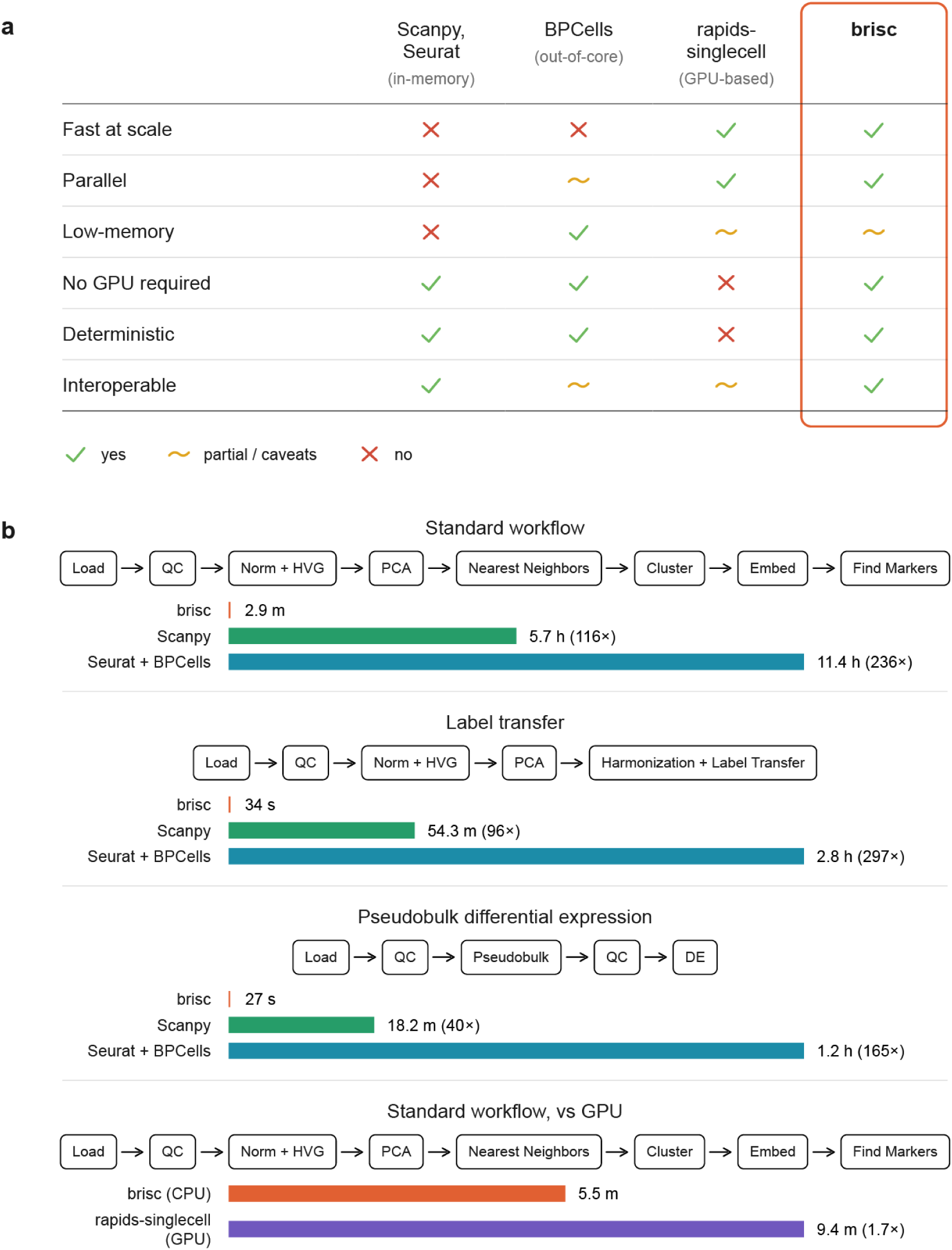
brisc analyzes millions of cells in minutes on CPUs. **a**, Strengths and limitations of the main strategies for large-scale single-cell analysis, each with a representative toolkit. brisc combines the speed of GPU toolkits with the determinism of CPU toolkits, without requiring a GPU. **b**, Runtime of three common workflows on the Parse dataset (9.7 million cells) versus Scanpy and Seurat on a 192-core CPU node, and of the standard workflow versus rapids-singlecell on a GPU node (96 cores, 4 NVIDIA H100 GPUs). Bars show the mean of three runs, labeled with the slowdown relative to brisc.

The second strategy, GPU acceleration, targets runtime. rapids-singlecell reimplements standard single-cell workflows on NVIDIA GPUs and reports hundreds of fold speedups over Scanpy^5^. However, GPU usage comes with unique challenges: GPUs are expensive and scarce, the NVIDIA software stack is difficult to install and maintain, and large datasets that cannot fit in the GPU’s memory incur large performance costs since they must be held partly in CPU memory or split across multiple GPUs. Most importantly, existing GPU toolkits do not give reproducible results: GPUs split computations across thousands of threads that execute in a different order on every run, leading to differences in floating-point roundoff that compound across the steps of a workflow^6^.

What if it were possible to accelerate single-cell analyses without having to contend with the limitations of out-of-core or GPU toolkits? The concept of a multithreaded CPU toolkit is particularly attractive in the single-cell space, since large datasets require computers with large amounts of memory, which tend to also come with lots of CPUs that are left idle by single-threaded toolkits. However, single-cell analyses are also particularly difficult to multithread relative to other genomic data modalities, since many of their most time-consuming steps (e.g. nearest-neighbor finding, clustering, embedding, integration) rely on finding interrelationships between cells, which makes them far from embarrassingly parallel. GPUs, with their tens of thousands of cores, can plow through difficult-to-parallelize problems through sheer brute force, but greater care and ingenuity is necessary to achieve more than modest speedups when parallelizing across mere dozens to hundreds of CPUs, especially under the constraint of exact reproducibility. This challenge has hindered the use of CPU-based multithreading in the single-cell space.

Here we present brisc, the first end-to-end multithreaded CPU toolkit for single-cell analysis. brisc spans preprocessing, dimensionality reduction, integration, label transfer, clustering, embedding, differential expression and plotting. It offers a strong guarantee of reproducibility: results must exactly match from run to run, even when varying the number of threads. brisc reads, writes and converts single-cell data in-memory between the AnnData, Seurat and SingleCellExperiment formats, so Python and R tools can be combined within a single analysis. Below, we describe brisc’s design and benchmark its speed, memory, accuracy and reproducibility against Scanpy, Seurat with BPCells and rapids-singlecell on datasets of up to 20 million cells.

## Results

### A multithreaded, deterministic and interoperable CPU toolkit

brisc aims to run as fast as possible on multi-core CPU systems. To achieve this, we reimplemented the core steps of standard single-cell workflows from scratch in Cython, which compiles to C++ and facilitates efficient parallel execution. Parallelization alone, however, accounts for only part of brisc’s speed: for many steps, brisc is several times faster than Scanpy and Seurat even on a single thread. brisc employs a toolbox of optimization strategies to ensure speed, memory efficiency and determinism (Fig. 2). These strategies are broadly applicable to scientific software in general, and we present them as lessons for anyone optimizing their own toolkit.

**Fig. 2.**
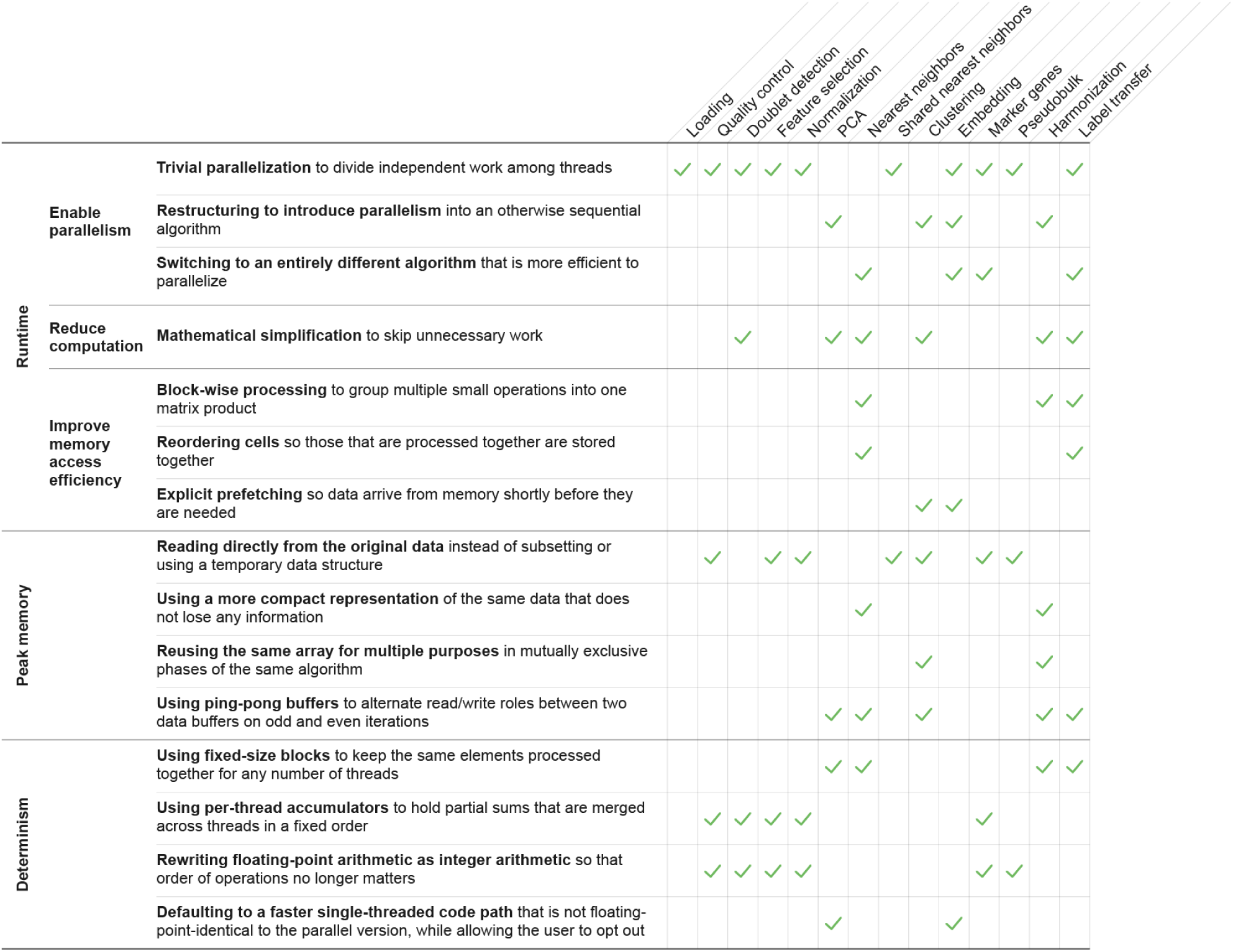
Optimization strategies behind brisc’s speed, memory efficiency and determinism. Optimization strategies (rows) grouped by whether they reduce runtime, reduce peak memory usage or ensure determinism, with checkmarks for the steps of brisc that use each one. Most strategies recur across several steps.

brisc reduces runtime by enabling parallelism, reducing computation, and improving memory access efficiency. Many steps, like normalization, parallelize trivially, but the slowest steps do not. For these, brisc either restructures the computation to expose parallelism or switches to an entirely different algorithm that is more amenable to parallelization. Where possible, brisc also reduces the number of mathematical operations by rewriting computations in mathematically equivalent ways.

For many steps, however, improving memory access efficiency matters far more for runtime than merely reducing computation. Modern CPUs have multiple levels of cache, which provide much faster access to data than main memory. Efficient use of this cache hierarchy is therefore critical for performance. brisc improves cache locality in two ways. First, it processes cells in blocks rather than individually, enabling highly efficient BLAS (Basic Linear Algebra Subprograms) matrix multiplications that reuse data while it remains in cache. Second, it reorders cells so that those processed together (e.g. nearest neighbors) are more likely to be located near each other in memory. This increases the likelihood that accessing one cell also brings nearby cells into cache, thereby speeding up subsequent accesses.

brisc further hides memory latency through explicit prefetching, which prompts the processor to load data into cache before it is needed. This is useful when the location of the next memory access depends on the result of the current one, making it difficult for the processor to predict automatically.

brisc takes special care to optimize memory access efficiency for the large fraction of multicore machines with non-uniform memory access (NUMA), where CPUs can access nearby memory regions faster than those attached to other CPUs. It uses a custom memory allocator to ensure each cell is placed in memory close to the CPU that will primarily process it.

These memory optimizations matter all the more because brisc is an in-memory toolkit. Unlike out-of-core toolkits like BPCells, which reread the data from disk at every step, brisc loads the dataset once and keeps it in memory. The trade-off is that the entire dataset has to fit in memory, and so brisc aggressively minimizes the amount of additional memory each step needs to use.

brisc is deterministic: every step is not only reproducible across repeated runs, but also gives identical results regardless of the number of threads used. This is unusual for parallel scientific software, where differences in thread scheduling can change the order of floating-point operations and thus the final result. brisc avoids this dependence on execution order, ensuring reproducible results without sacrificing parallelism (Fig. 2). There are two exceptions to this guarantee. First, the parallel implementations of principal component analysis (PCA) and of the embedding methods Pairwise Controlled Manifold Approximation (PaCMAP)^7^ and its Local Adjusted Graph variant (LocalMAP)^8^ incur overhead that is unnecessary for single-threaded execution. brisc therefore defaults to faster single-threaded implementations that are reproducible across runs but not bitwise-identical to their parallel counterparts. Users can opt out of these implementations if they wish to obtain identical results to the parallel versions. Second, because Uniform Manifold Approximation and Projection (UMAP)^9^ cannot be efficiently parallelized while preserving determinism, brisc offers a nondeterministic parallel implementation that users must explicitly enable.

brisc is also uniquely interoperable with existing single-cell ecosystems. It loads and saves the file formats used by Scanpy, Seurat and Bioconductor, as well as raw 10x Genomics data. It also converts datasets in-memory to and from AnnData, Seurat and SingleCellExperiment objects, without writing to disk, so analyses can move seamlessly between brisc, these other toolkits, and packages in their ecosystems – even those written in R – within the same pipeline. Even bespoke analyses that brisc does not directly implement can still benefit from brisc’s parallelism, since brisc also parallelizes basic operations like subsetting, subsampling and splitting by cell type, which are often the slow parts of custom analyses. brisc is also cross-platform, and is available for Linux, Mac, and Windows.

Finally, brisc checks rigorously for errors. Much of what makes single-cell analysis hard for beginners is that mistakes often produce hard-to-understand errors deep inside a toolkit, forcing the user to understand low-level implementation details in order to fix them. Worse, some common mistakes, like passing normalized counts to a step that needs raw counts, do not give an error at all, but still silently produce incorrect results. brisc strictly validates the types of its inputs and explicitly tests for these sorts of common mistakes, providing concise but actionable error messages that explain not just what went wrong but also how to fix it.

### Performance benchmarks

We benchmarked brisc against Scanpy, Seurat and rapids-singlecell on three public datasets: the Seattle Alzheimer’s Disease Brain Cell Atlas (SEA-AD) of 1.4 million human brain cells^10^; a Parse Biosciences dataset of 9.7 million peripheral blood mononuclear cells (PBMCs)^11^; and the PanSci mouse atlas of 20.3 million cells from 13 organs^12^. Since Seurat can only analyze datasets with up to 2^31^ − 1 (∼2.1 billion) non-zero counts, and all three datasets are several times larger than that, we paired Seurat with BPCells, which streams the count matrix from disk and is what Seurat’s developers recommend for large datasets.

We benchmarked three workflows (Fig. 1b): a standard workflow encompassing cell-type clustering, embedding, and marker gene detection (Fig. 3 and Extended Data Fig. 1); integration and label transfer from an annotated reference (Extended Data Fig. 2); and differential expression (Extended Data Fig. 3). In addition to these three workflows, we also benchmarked 6 common data manipulation operations (Extended Data Fig. 4). We ran each workflow three times, both to measure variability in runtime across runs and to assess reproducibility. We benchmarked the three CPU toolkits on a node with 192 cores and 749 GB of memory, and separately benchmarked brisc against rapids-singlecell on the standard workflow on a node with 96 cores and 4 NVIDIA H100 GPUs.

**Fig. 3.**
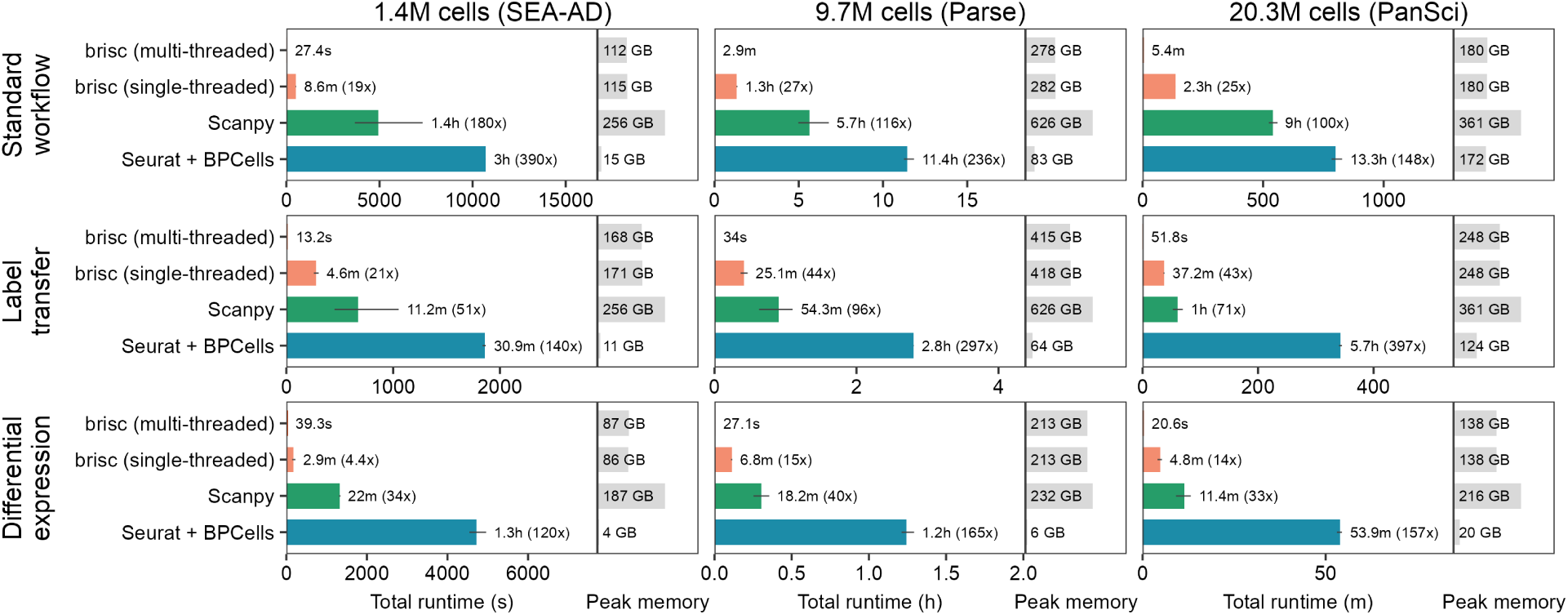
Runtime and peak memory of three analysis workflows across toolkits and dataset sizes. Total runtime and peak memory usage of three analysis workflows (rows) on three datasets (columns), run on a 192-core CPU node. brisc was run two ways, using all cores (multi-threaded) and using one core (single-threaded). Bars show the mean runtime of three runs and the slowdown relative to multi-threaded brisc; error bars show the range. Gray bars show peak memory, the maximum across the three runs. Runtime axes are scaled independently for each panel and use different units for each dataset.

brisc ran the standard workflow in 27 seconds on SEA-AD, 3 minutes on Parse and 5.4 minutes on PanSci; Scanpy took 100–180× longer and Seurat took 148–390× longer (Figs. 1b and 3). Even single-threaded, brisc was 4–10× faster than Scanpy. Label transfer and differential expression took brisc under 55 seconds on every dataset; Scanpy took 33–96× longer and Seurat 120–397× longer. Despite holding the entire dataset in memory, brisc’s peak memory usage on the standard workflow was about half of Scanpy’s on every dataset (Fig. 3). Seurat, which streams from disk, used 3–8× less memory than brisc on SEA-AD and Parse, but about as much as brisc on PanSci. We discuss how brisc achieves such favorable results in the sections below, and provide per-step benchmarks in Fig. 4 and Extended Data Figs. 1–3.

**Fig. 4.**
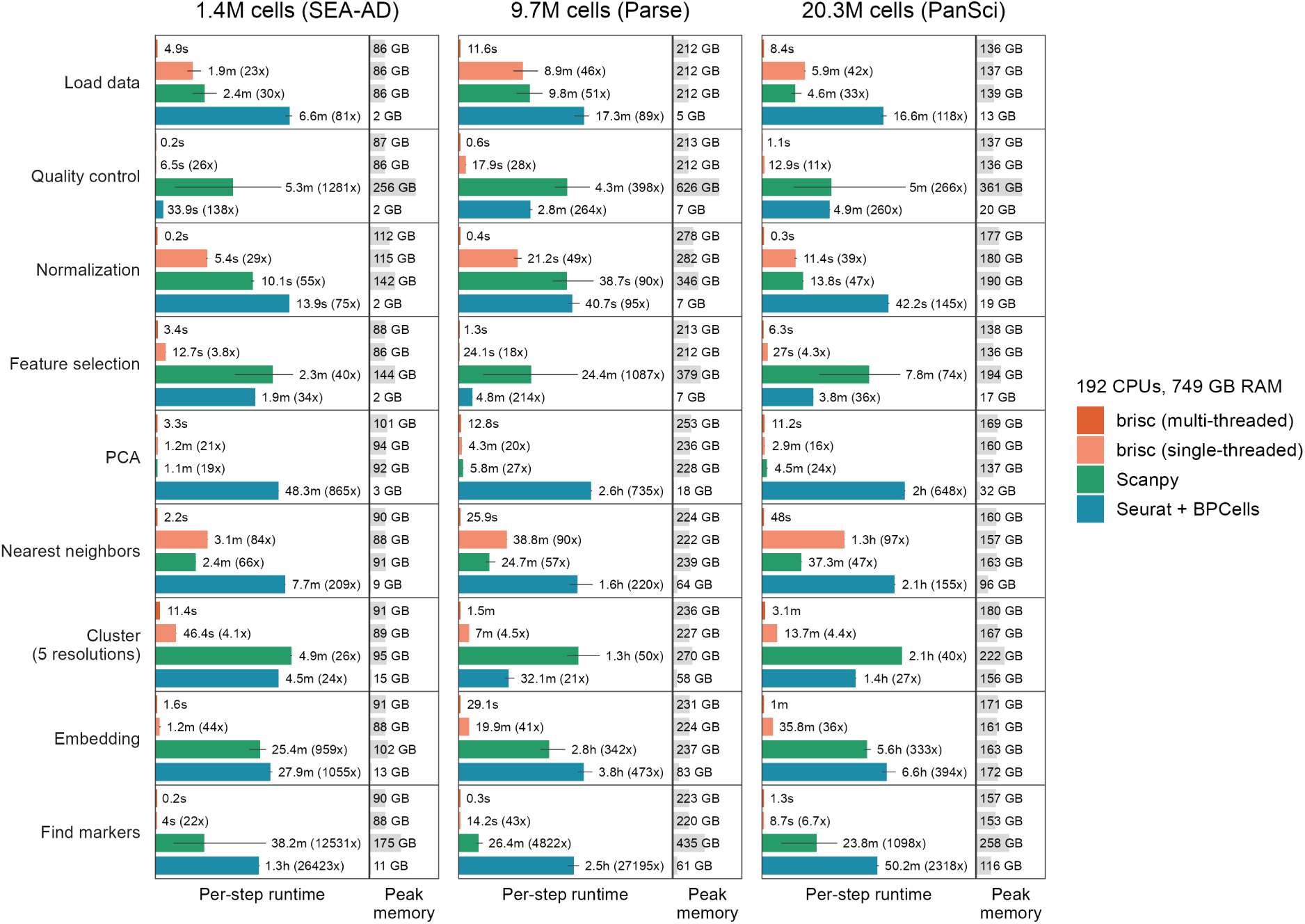
Runtime and peak memory usage of each step of the standard workflow. Runtime and peak memory of each step of the standard workflow (rows) on three datasets (columns), from the same runs as in Fig. 3. Display conventions as in Fig. 3. Seurat’s “Load data” entry is the time taken to convert the input file to the on-disk BPCells format. brisc computes the embedding with LocalMAP and the other toolkits with UMAP.

### Data loading

A substantial bottleneck in large-scale single-cell analysis is data loading, because the HDF5 library that all major toolkits use to read .h5ad files (libhdf5) does not support multithreaded reading. brisc gets around this in two ways. For uncompressed and unchunked HDF5 files, including those in our benchmarks, it bypasses libhdf5 entirely and implements a custom multithreaded reader, which on Linux networked file systems improves performance several-fold by bypassing the operating system’s page cache. For files with compression or chunking, it spawns multiple processes that each read part of the file into shared memory (Methods). In our benchmarks, brisc loaded all three datasets in under 12 seconds (Fig. 4), 30– 51× faster than Scanpy and 81–118× faster than Seurat.

### Preprocessing and dimensionality reduction

Quality control (QC), feature selection and normalization took a total of under 10 seconds on all three datasets (Fig. 4), 101–727× faster than Scanpy and 43–206× faster than Seurat. Feature selection and normalization operate on each gene or cell independently, so brisc simply parallelizes them. For QC, brisc avoids subsetting the dataset to QC-passing cells, instead keeping track of them with a Boolean array that downstream steps use to decide which cells to skip. This strategy avoids having the original and QCed data briefly coexist in memory, which nearly doubles peak memory usage and led to QC being Scanpy’s most memory-intensive step on all three datasets (Fig. 4). Similarly, Scanpy subsets the count matrix for each batch when selecting highly variable genes within batches, which incurs runtime and memory overhead, whereas brisc does not.

PCA is harder to parallelize. brisc uses the same approximate PCA algorithm as Seurat, which spends almost all its time multiplying the sparse count matrix and its transpose by a vector. Sparse matrix-vector products only parallelize efficiently when the sparse matrix is in row-major format rather than column-major format, and since the transpose of one is the other, half of the products are stuck single-threaded. To circumvent this limitation, brisc creates both a row-major and a column-major copy of the count matrix subset to highly variable genes, so that both types of products can run efficiently in parallel (Methods). This is still reasonably memory-efficient because only the highly variable genes are copied. brisc computed principal components in under 13 seconds across all three datasets, 19–27× faster than Scanpy and 648– 865× faster than Seurat.

### Nearest neighbors

brisc’s approximate nearest-neighbors search was both the fastest and the most accurate of the four toolkits’. On all three datasets, brisc found each cell’s nearest neighbors and constructed the shared nearest neighbor graph in under 50 seconds, 47–66× faster than Scanpy and 155–220× faster than Seurat (Fig. 4). brisc made 82% fewer errors than Seurat and 98% fewer than Scanpy on SEA-AD, 81% fewer than Seurat and 94% fewer than Scanpy on Parse, and 81% fewer than Seurat and 97% fewer than Scanpy on PanSci. On Parse, the dataset all toolkits performed worst on, brisc still recovered 99% of each cell’s true neighbors, compared with 95% for Seurat and rapids-singlecell and 86% for Scanpy (Fig. 5). The gap was larger in the quarter of cells where each toolkit performed worst, for which brisc still recovered 96% of true neighbors, compared with 84% for Seurat and rapids-singlecell and 65% for Scanpy (Fig. 5).

**Fig. 5.**
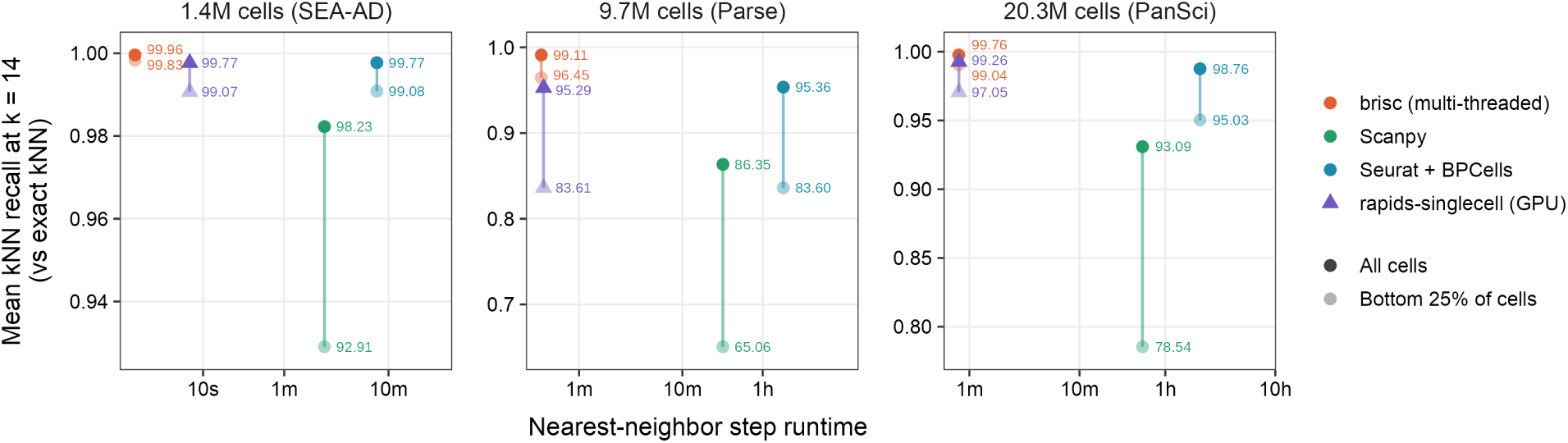
Accuracy and runtime of the nearest-neighbor search. Mean recall of each toolkit’s approximate nearest-neighbor search against exact nearest neighbors, computed by brute force from that toolkit’s principal components (k = 14, the largest neighbor count returned by every toolkit at default settings), plotted against the mean runtime of the nearest-neighbor step (three runs, as in Fig. 4). Dark points show the mean over all cells and light points show the mean over the 25% of cells with the lowest recall. rapids-singlecell (triangles) ran on the GPU node and the other toolkits on the CPU node. Runtime axes are log-scaled and independent for each dataset.

brisc’s search follows the inverted file index strategy of the Facebook AI Similarity Search (FAISS) library^13^. It first clusters cells with k-means, then exhaustively searches each cell’s 64 nearest clusters, so a true neighbor is missed only when it lies outside those clusters. brisc adds three interlinked optimizations that reduce how often cells are loaded from memory (Methods). First, brisc pre-sorts the cells so that they are grouped by cluster, with the clusters ordered according to their centroid’s first principal component, so that cells near each other in principal component space are also near each other in memory. Second, brisc searches consecutive cells together in blocks, so each cluster is loaded from memory once per block rather than once per cell. Third, brisc computes block-to-cluster distances as a single, highly efficient matrix multiplication, leveraging a mathematical simplification that removes a part of the distance calculation that is constant. Together, these optimizations make brisc’s search about an order of magnitude faster than FAISS’s, without changing the result.

### Clustering

Clustering is one of the slowest steps of the standard workflow, and is generally rerun at several resolutions. brisc retains the standard Leiden algorithm but reimplements it from the ground up. Even on a single thread, brisc’s Leiden clustering is 5–11× faster than the igraph library^14^ that Scanpy uses. This speedup is entirely due to systems engineering. brisc uses a more compact layout of the data in memory, allocates memory once rather than repeatedly, prefetches data from memory several steps ahead of when it is needed to hide memory latency, and performs less arithmetic in the innermost steps (Methods). Since Leiden clustering cannot be efficiently parallelized within a single resolution without giving up determinism, brisc parallelizes across resolutions instead. Clustering was brisc’s slowest step on all three datasets: brisc clustered SEA-AD at five resolutions in 11.4 seconds, Parse in 1.5 minutes, and PanSci in 3.1 minutes (Fig. 4). Across the three datasets, Scanpy took 26–50× longer and Seurat 21–27× longer. brisc’s clusters had similar or better agreement with the original datasets’ cell-type annotations than the other toolkits’ clusters (Extended Data Fig. 5).

### Embedding

brisc supports three methods for embedding cells in two dimensions for visualization. UMAP, the standard approach, preserves local structure well but tends to distort the global arrangement of clusters. PaCMAP, a newer method, preserves both local and global structure, and LocalMAP is a variant of PaCMAP that aims to prevent distinct clusters from collapsing into one another. We benchmarked brisc with LocalMAP (see Extended Data Fig. 6 for all embeddings), which unlike UMAP can be parallelized deterministically. brisc took under 75 seconds on all three datasets (Fig. 4), 333–959× faster than Scanpy and 394–1,055× faster than Seurat. UMAP was the slowest step of the standard workflow in Scanpy and Seurat for Parse and PanSci, taking several hours on each. Seurat also reached its peak memory usage at the embedding step on Parse and PanSci, because its UMAP runs entirely in memory.

We also benchmarked brisc’s UMAP implementation against Scanpy and Seurat. brisc’s implementation is based on the original umap-learn implementation of UMAP (also used by Scanpy), but uses a rejection sampling-style approach rather than deterministic scheduling to decide which pairs of cells have their positions adjusted on each iteration of the layout optimization. This approach reduces memory traffic by avoiding the need to repeatedly access scheduling metadata for every pair of cells. Single-threaded, brisc’s UMAP was about 4× faster than Scanpy’s and 5× faster than Seurat’s (Extended Data Fig. 6). Unlike Scanpy and Seurat, brisc also supports running UMAP in parallel via the Hogwild! algorithm^15^, which is 11– 18× faster than Scanpy and 13–21× faster than Seurat across the three datasets. Even multithreaded, brisc’s UMAP was still over 24× slower than PaCMAP and over 22× slower than LocalMAP on every dataset, underscoring the runtime benefits of these alternative embedding methods (Extended Data Fig. 6).

### Marker genes

brisc finds marker genes with a simpler and faster approach than conventional tools. Whereas Scanpy and Seurat test every gene for differential expression between cell types, brisc instead selects marker genes based on their detection rate (fraction of cells with non-zero expression) in each cell type: it should be high in the target cell type and many times smaller in other cell types^16^. These two criteria trade off against each other, since genes with high detection rates in one cell type tend to be more broadly expressed in other cell types. As a result, brisc selects genes on the Pareto front of the two metrics: those that no other gene beats on both metrics simultaneously. Because detection rates are so cheap to compute, this approach is extremely fast, taking under 2 seconds on all three datasets, 1,100–13,000× faster than Scanpy and 2,300– 27,000× faster than Seurat.

### Label transfer

Harmonization removes batch effects so that cells group by cell type rather than by dataset. Once a new dataset has been harmonized with a well-annotated reference, label transfer annotates it by giving each cell the most common label among its nearest reference cells. Transferring labels from a reference is increasingly used as an alternative to manual annotation of clusters based on marker genes. Mapping a dataset to several references and comparing the results is more informative than relying on one. In our experience, users value brisc most for its speed at these two steps, which makes trying several references easy.

brisc includes a multithreaded reformulation of version 2 of the widely used Harmony algorithm^17^. Harmony is hard to parallelize because of a constraint built into the algorithm. It works through the cells in a sequence of blocks, and each block update depends on statistics accumulated over all the blocks before it, so no two blocks can be processed in parallel without changing the result. As a result, the reference R implementation has no parallel code of its own: while it allows setting the number of threads, this only affects BLAS calls, which are a negligible fraction of total runtime.

brisc’s innovation is to parallelize Harmony within each block, by dividing each block into fixed-size chunks which are processed in parallel. The result is identical regardless of thread count. brisc also simplifies Harmony by representing batch membership as a vector instead of a “one-hot” matrix (which reduces memory traffic and enables a variety of mathematical simplifications), and by computing Harmony’s objective function only when needed, rather than after every clustering iteration.

brisc’s label transfer uses nearest-neighbor search to find each query cell’s closest cells in the reference. Whichever cell-type label is most common among these closest cells, brisc assigns to the query cell. brisc also reports the percent of these closest cells that agree on the cell-type label, as a metric of confidence in the cell-type call. Harmonization and label transfer together took brisc under 30 seconds on all three datasets (Extended Data Fig. 2), 90–156× faster than Scanpy’s ingest method and 428–972× faster than Seurat’s anchor-based approach. brisc labeled 87–99% of query cells correctly (relative to the cell-type calls that came with the dataset) across the three datasets, as accurately as Seurat and Scanpy, except on SEA-AD, where Scanpy labeled only 74% correctly (Extended Data Fig. 7).

### Differential expression

brisc uses pseudobulk limma-voom for differential expression testing. Pseudobulking took brisc about a second or less on every dataset, whereas Scanpy and Seurat took up to half a minute. Differential expression testing took brisc under 35 seconds on every dataset (Extended Data Fig. 3), and across the three datasets Scanpy took 34–35× longer and Seurat 133–253× longer (Methods).

brisc’s differential expression framework is dramatically simpler and more flexible to use than other toolkits’. The user specifies an R formula, such as ∼ disease_status + age + sex, and a single function call runs the full limma-voom pipeline across every cell type; custom contrasts can also be specified. This is simpler than even running limma-voom directly in R, which takes several steps. brisc gives each of these steps a sensible default, and any of them can be overridden, per cell type if needed. Whenever the comparison is between groups, such as cases versus controls, brisc uses voomByGroup^18^ rather than plain voom, which fits a separate mean-variance trend for each group to improve calibration. Due to the large number of use-cases supported, it is also the most extensively error-checked function in brisc, incorporating detailed quality control and input validation at each step.

### Data manipulation

brisc also accelerates the everyday operations that fall between the steps of a workflow, like subsetting a dataset or splitting it by cell type. Custom analyses are full of these operations, and because other toolkits do not optimize them much, they are often the slow part. This is especially true of out-of-core toolkits, since any operation the toolkit does not explicitly implement requires loading the entire count matrix into memory, which is exactly what out-of-core computation exists to avoid. brisc extends SciPy’s sparse matrix data type so that expensive subsetting and reformatting operations on the count matrix automatically run in parallel, and stores cell and gene metadata in polars data frames, which also automatically parallelize many operations.

We timed six everyday operations: looking up a single gene’s expression, subsetting to one cell type, subsetting to highly variable genes, subsampling, splitting by cell type, concatenating cell types after they have been split. brisc took under 4 seconds for each of these operations on every dataset, whereas Scanpy took 5 minutes to split Parse by cell type and Seurat took 23 minutes to concatenate PanSci’s cell types (Extended Data Fig. 4).

### Comparison with GPUs

On SEA-AD, brisc completed the standard workflow in 46 seconds, compared with 3.8 minutes for rapids-singlecell; on Parse, in 5.5 minutes compared with 9.4 minutes; and on PanSci, in 10 minutes compared with 8.5 minutes (Fig. 1b and Extended Data Fig. 1). Each dataset has several times more non-zero counts than a GPU sparse matrix can hold, and all three exceed the memory of a single GPU, so we were forced to run rapids-singlecell in its Dask-based multi-GPU mode, with each dataset split into chunks across the four GPUs (Methods).

### Reproducibility

Finally, we tested whether each toolkit gives the same result when the same analysis is rerun. On each dataset, we compared the output of each step of the standard workflow between the three runs (Fig. 6 and Methods). brisc, Scanpy and Seurat always gave identical results, but rapids-singlecell’s output changed between runs, and the differences grew at each step. Its quality control and feature selection were identical, but its principal components differed slightly, with a root-mean-square difference of at most 1.9 × 10^−4^ standard deviations per component. That was enough to change 0.3–6.0% of each cell’s nearest neighbors. By clustering, 15–26% of cells changed cluster membership between runs, and the three runs disagreed on the number of clusters on Parse (finding 32, 35, or 37 clusters depending on the run) and PanSci (finding 55, 56, or 59).

**Fig. 6.**
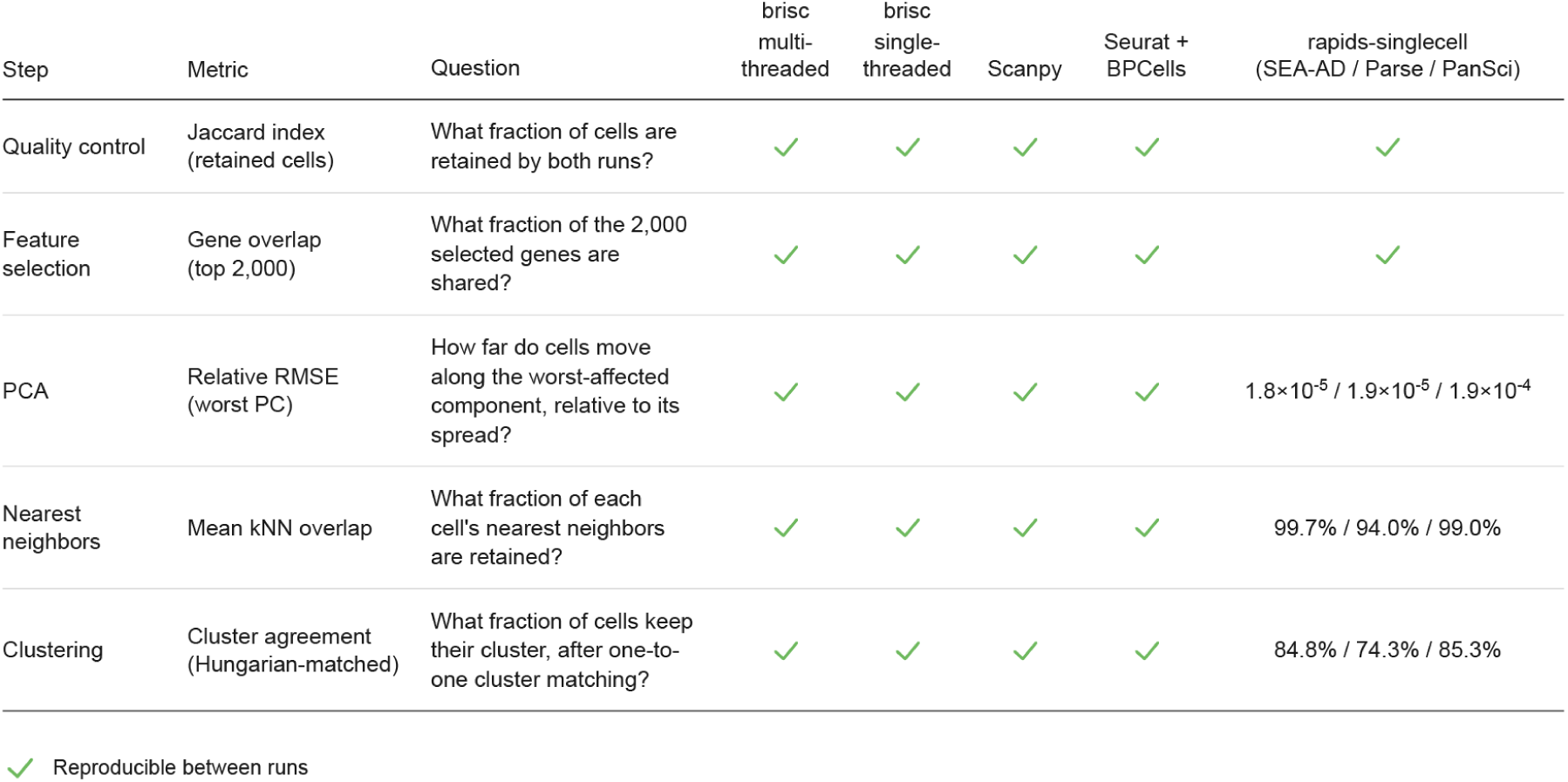
Run-to-run reproducibility of steps of the standard workflow. Agreement of each step’s output between each pair of the three runs of the standard workflow, for each toolkit and dataset. Checkmarks indicate identical output in all three runs. Where output differed, the entry gives the worst of the three pairwise comparisons on each dataset (SEA-AD / Parse / PanSci). Clustering is compared at a resolution of 1. Only rapids-singlecell differed between runs, starting with small differences in the principal components that grew at each later step.

## Discussion

brisc is the first general-purpose multithreaded CPU toolkit for single-cell analysis, and it allows the analysis of tens of millions of cells to become near-real-time. In practice, this expands the space of analysis choices and hypotheses that can be tested within a given period of time. A user can load a raw dataset, process it fully and examine the clusters, markers and embedding in minutes rather than waiting until the next day. They can then revise analysis parameters like quality control filters and see the effect in a few minutes more. brisc’s speed is particularly useful for analyses that span many datasets, like harmonizing several large datasets to explore whether they form a coherent atlas, transferring labels from multiple reference datasets and comparing the results, and meta-analyzing differential expression across several independent datasets for the same disease. Production pipelines benefit as well, since cell atlas consortia, core facilities and commercial platforms run the same steps on every dataset they process and need both throughput and reproducible results. Given how much faster brisc is than existing toolkits, we expect it to find uses and lead to analysis strategies that we have not anticipated.

More broadly, brisc demonstrates how much faster software for biology can be. As speed becomes easier to achieve, good software will be distinguished from merely fast software by two properties that brisc was designed around. The first is reproducibility. Developers and users of parallel biology software often tolerate results that vary slightly between runs, but brisc makes strict determinism a core design principle, enabling efficient and reproducible science. On the rare occasion that a step cannot be efficiently parallelized without losing determinism – which in brisc only occurs in UMAP – non-determinism is permissible, but the user must explicitly opt into it. The second property is rigorous input checking: brisc validates its inputs before doing any substantial work, and when an input is invalid, it says what is wrong and how to fix it. We present brisc as a template for how to build fast, reproducible, and user-friendly software in other areas of bioinformatics.

The GPU toolkit we benchmarked against is a cautionary tale. rapids-singlecell on four H100 GPUs – a more powerful GPU setup than many users would have access to – was comparable in speed to brisc on 96 CPUs, but its results differed on every run. The differences began as rounding differences in the principal components and ended with over a quarter of cells changing their cluster membership, and on the two largest datasets, no two runs agreed on the number of clusters. This is not a cosmetic discrepancy; it is a fundamental flaw. Clusters define the units of downstream biological interpretation, like which cell types and marker genes are identified, and which gene expression changes are deemed significant. When over a quarter of cells change cluster assignment, some biological conclusions will change.

There is also a practical argument for using CPUs instead of GPUs to achieve parallelism. GPUs are expensive and scarce due to their popularity for AI applications, and there are no clear signs of this easing within the next few years (though longer-term trends in this fast-moving space are hard to predict). Every analyst already has multicore CPUs in their laptops, and software that uses them well democratizes science in a way that GPU software cannot. This matters most for users in low-and middle-income countries (LMICs), where a state-of-the-art GPU server may be out of reach, but a multicore workstation is not.

Out-of-core computation, the other strategy for large datasets, is in our view the wrong foundation for a single-cell toolkit, due to its lack of interoperability with third-party tools. Streaming the count matrix from disk saves memory only for steps that are explicitly designed to use it. For instance, even interfacing BPCells with Seurat in the way Seurat’s developers recommend led the embedding step to run entirely in-memory, with peak memory usage nearly matching brisc’s on PanSci’s 20 million cells. Even so, Seurat with BPCells was the most memory-efficient toolkit overall: for instance, brisc required 278 GB of memory and Scanpy required 626 GB to analyze Parse, the most memory-hungry of the three datasets for these two toolkits, while Seurat with BPCells required only 83 GB. In the near term, memory constraints will tighten, because datasets are growing faster than the amount of memory available on new computers, and demand from AI has halted the decades-long decline in memory prices. In the longer term, however, we expect memory constraints to ease. Storage that is persistent like a disk but even faster than current RAM already exists in prototype^19^, and widespread use of such storage would remove the need to compute out-of-core at all.

brisc currently analyzes only single-cell and single-nucleus transcriptomics data, and extending it to multimodal data is the main direction for future versions. Pairing transcriptomics with measurements of a cell’s spatial location (spatial transcriptomics), chromatin accessibility (multiome) or cell-surface protein markers (CITE-seq) would add new data types and analysis steps, but the core of brisc would apply to these modalities unchanged, so supporting them is comparatively little extra work. With these additions, we expect fast, reproducible analysis of tens of millions of cells to extend to every kind of single-cell data.

## Methods

### Core design principles Fast and parallel

brisc is a parallel CPU-based toolkit and aims, above all, to run as fast as possible on large multi-core machines. It reimplements core steps of standard single-cell workflows from the ground up, using algorithms and data structures designed for parallel execution and cache-efficient memory access. Most steps make extensive use of Cython, which compiles to optimized C++ code and enables efficient multithreading by circumventing Python’s global interpreter lock (GIL).

brisc’s parallelism is built into the core data structures it uses to represent single-cell data. To accelerate sparse matrix operations, like subsetting and format conversion of the count matrix, brisc implements a custom multithreaded subclass of scipy’s sparse arrays that is fully backward-compatible with scipy. For per-cell and per-gene metadata, brisc uses the Rust-based polars data frame library, which automatically parallelizes operations across columns.

brisc is designed to perform well even on non-uniform memory access (NUMA) Linux systems, where different CPU cores have different speeds of access to different regions of memory. Such systems are increasingly the norm for machines with large numbers of CPUs, such as those in high-performance computing clusters. brisc features a custom NUMA-aware memory allocator that bypasses glibc’s malloc arena to ensure memory is allocated close to the CPU that will use it (strict first-touch placement), improving memory access speeds and reducing runtime.

### In-memory

brisc is an in-memory toolkit. Unlike “out-of-core” toolkits like BPCells^3^ and Scarf^4^ that repeatedly load chunks of the data from disk every time they need to be used, brisc loads the entire dataset once and keeps it in memory for all subsequent operations. This dramatically reduces I/O traffic and improves performance, particularly on the distributed file systems that are standard on large-scale computing clusters. The tradeoff of this approach is that it is substantially more memory-hungry than out-of-core approaches. brisc mitigates this by aggressively memory-optimizing each step through a variety of strategies, including avoiding temporary copies or data structures, using more compact data representations, reusing buffers during mutually exclusive phases of the same algorithm, and using ping-pong buffers (Fig. 2). In practice, on large datasets, brisc has about half the peak memory usage of Scanpy, which is itself more memory-efficient than Seurat for in-memory analyses.

### Deterministic

brisc is a fully deterministic toolkit. Every step (except multi-threaded UMAP, discussed below) is reproducible across repeated runs, but also gives identical results regardless of thread count. (For single-threaded PCA, PaCMAP, and LocalMAP, identical results to the parallel version must be explicitly enabled with match_parallel=True, as discussed below.) Both properties are highly unusual for parallel scientific software. It also gives identical results (for PCA, with match_parallel=True only) regardless of whether the count matrix is stored in row-major (also known as compressed sparse row) or column-major (compressed sparse column) format.

Irreproducibility across repeated runs (non-determinism) occurs because threads are not guaranteed to execute in the same order, and this can affect the order of floating-point operations. Floating-point arithmetic is not associative – (A + B) + C is not guaranteed to equal A + (B + C) – so different orders of execution can produce different results, even when the underlying math is the same. Even if a program is made deterministic for a fixed number of threads, changing the thread count typically changes how work is partitioned across threads, again altering the order of floating-point operations and producing divergent results.

brisc uses a variety of strategies to avoid both pitfalls. To eliminate run-to-run irreproducibility, brisc avoids synchronization between threads (it is often highly inefficient in any case). To additionally ensure consistency across thread counts, when performing floating-point reductions (i.e. summing multiple array elements into a scalar), brisc has each thread operate on fixed-size blocks of the array, so that block boundaries are independent of thread count. Other strategies brisc uses include creating per-thread buffers that hold partial sums, which are then merged serially across threads in a fixed order; rewriting floating-point arithmetic as integer arithmetic where possible, so that the order of operations does not affect the results; and providing a faster single-threaded code path that is used by default, while allowing the user to opt into exact floating-point identity with the parallel version (Fig. 2). Certain steps use additional, bespoke parallelization strategies, such as our “dual-matrix” strategy used during principal component analysis (see below).

### Interoperable

Uniquely among single-cell toolkits, brisc can directly load, save, and convert in-memory to and from datasets from all three major single-cell ecosystems – scverse/Scanpy (.h5ad files and in-memory AnnData objects), Seurat (.rds and .h5Seurat files and in-memory Seurat objects), and Bioconductor (.rds files and in-memory SingleCellExperiment objects) – as well as raw 10x Genomics data (.h5 or .mtx/.mtx.gz). For .h5Seurat and .h5 files, which are based on the same HDF5 file format as .h5ad files, brisc uses the same optimized parallel HDF5 reader as for .h5ad files. For .rds files, brisc uses R for reading and writing, leveraging the ryp Python-R bridge (https://github.com/briscverse/ryp) to quickly convert data between Python and R in-memory. Crucially, conversion to and from AnnData objects is zero-copy for the count matrix and largely zero-copy for the metadata as well, and therefore occurs almost instantaneously, with almost no memory overhead. This allows even the largest datasets to interface seamlessly with Scanpy and third-party libraries. Conversion to R is limited by the fact that the sparse matrix class used by Seurat and SingleCellExperiment, dgCMatrix, cannot represent single-cell datasets with more than 2,147,483,647 (INT32_MAX) non-zero elements – which, for reference, is smaller than all three datasets we use for benchmarking – but can nonetheless be done largely zero-copy. brisc’s interoperability makes it possible to seamlessly interleave Python and R workflows within the same analysis pipeline, without the I/O overhead of writing intermediate files to disk or the computational and memory overhead of repeated data copies.

### User-friendly

brisc is designed with user-friendliness as a top priority alongside efficiency. It features sensible defaults, strict type-checking, and solution-focused error messages. brisc explicitly checks for common beginner errors, like running steps before their prerequisite steps have been run or using normalized data for a step that requires unnormalized data (or vice versa). brisc largely avoids in-place operations, which are prone to accidentally overwriting data, and requires the user to specify overwrite=True for operations that would overwrite an existing data field or a file on disk. Because each step returns a new dataset, workflows can be written concisely using method chaining syntax (e.g. sc.qc().hvg().normalize().pca()).

### Data loading

A substantial bottleneck in large-scale single-cell analysis is data loading, which standard toolkits perform single-threaded. This is because the core software for reading the major single-cell data formats was not designed for parallelism. For instance, several major formats (.h5ad, .h5, and .h5Seurat) are built on HDF5, a general-purpose container for large arrays. All toolkits’ HDF5 readers rely on the same underlying library, libhdf5, which uses locks to restrict reading to one thread at a time.

The only reliable way to circumvent libhdf5’s locking is to use multiprocessing instead of multithreading, but this raises the issue of how to efficiently transmit the data back to the parent process after reading. To address this challenge, brisc allocates shared-memory buffers in the parent process, and has worker processes read directly into them, avoiding intermediate copies or additional memory usage. Each worker reads non-overlapping chunks of the input arrays, so no synchronization is necessary.

When reading HDF5 files saved without compression or chunking, brisc avoids using libhdf5 to read the count matrix and cell and gene metadata, instead opting to directly read the file’s raw bytes into memory via the pread() system call on Linux and Mac or the ReadFile() system call on Windows. String datasets (like cell and gene names) require special handling because they are stored non-contiguously within the file, but can still be read in parallel via indirect lookups. This approach is not dramatically faster than the multiprocessing-based reader when reading from local filesystems (since the bottleneck is I/O rather than process-spawning or shared-memory overhead) but offers several-fold speedups on Network File Systems (NFS) because brisc asks the operating system to bypass the page cache during reading (using the O_DIRECT flag to the open() system call on Linux and the F_NOCACHE flag to the fcntl() system call on Mac) on filesystems that support this.

### Quality control

The first step in most single-cell workflows is quality control (QC). At a high level, single-cell QC aims to remove doublets (a “cell” that is actually two cells sharing the same droplet), empty droplets (or droplets that contain only cell fragments), and damaged cells. Conventionally, empty droplets and damaged cells are identified based on having a high proportion of mitochondrial reads (e.g. >5%) and/or too few genes with non-zero expression (e.g. <100), while a variety of algorithms have been developed for doublet detection. brisc follows the same general approach, filtering to cells with ≤5% mitochondrial reads and ≥100 genes detected, along with optional doublet detection (see next subsection). A unique feature of brisc’s quality control is that it also filters to cells with non-zero ***MALAT1*** expression: ***MALAT1*** is a nuclear-expressed lncRNA, and cells with zero expression likely represent empty droplets or poor-quality cells^20^.

The most notable feature of brisc’s QC implementation is that by default, it avoids subsetting the dataset to cells passing QC, instead maintaining a Boolean mask of QC-passing cells. Downstream steps read this mask when deciding which cells to operate on. Since most cells in typical single-cell datasets pass QC, standard single-cell pipelines must temporarily store both the original count matrix and a QCed count matrix of almost the same size in memory. This means that avoiding subsetting roughly halves the peak memory usage of the QC step, which would otherwise be the memory bottleneck for most standard single-cell workflows.

### Doublet detection

brisc uses an efficient parallel implementation of coexpression-based doublet scoring (cxds)^21^, one of the fastest doublet-detection algorithms. Since the cxds score is continuous, it needs to be converted into a binary classification of doublets versus non-doublets. This problem can be framed as finding a cxds score threshold above which a cell is deemed to be a doublet. To determine this threshold, brisc simulates doublets by combining the counts from randomly selected pairs of cells, via the following steps:

1. Sample as many random pairs of cells (with replacement) as there are real cells.
2. Combine the counts from each pair of cells into a simulated doublet. Because cxds operates on binarized count matrices, brisc averages the two cells’ count matrices in a binary sense: if a gene is expressed in either cell, it is deemed to be expressed in the simulated doublet, but if it has a count of 1 in one cell and 0 in the other, it is randomly chosen to be either expressed or not expressed with equal probability (since the average count would be 0.5).
3. Calculate cxds scores for these simulated doublets, based on the coexpression patterns (the S matrix from cxds) learned from the real data.
4. Take the median cxds score of the simulated doublets as the threshold. In other words, if a real cell has a higher doublet score than the average simulated doublet, brisc calls it a doublet.

Since doublets cannot occur across batches, brisc allows the user to specify each cell’s batch, and performs doublet detection separately for each batch.

### Highly-variable gene identification

brisc identifies highly-variable genes via a parallel implementation of the algorithm developed by Seurat in FindVariableFeatures() and implemented in Scanpy as the seurat_v3 flavor of highly_variable_genes(). However, brisc’s implementation offers three key improvements over conventional implementations. First, it uses integer summation when calculating the sums and squared sums that go into the variance calculation, deferring the use of floating-point arithmetic until the final variance calculation. This reduces floating-point error on datasets with millions of cells. Second, when running multi-batch highly variable gene selection, which is increasingly the norm, brisc avoids subsetting the count matrix for each batch, reducing runtime and memory requirements. Third, for row-major count matrices, brisc avoids most of the work in the second pass over the count matrix, which calculates each gene’s variance after clipping its counts at a gene-specific maximum. Because clipping affects only rare, extreme counts, brisc derives the clipped sums from the unclipped sums calculated in the first pass, correcting them for only the counts above the clipping threshold.

### Normalization

brisc supports three normalization methods. All three methods normalize each cell independently of the rest and log-transform the counts in some way, but differ in the details.

The simplest approach (“logCP10k”) computes the natural logarithm of the number of counts that belong to a gene per 10,000 total counts in its cell, plus one: log(count / total_counts × 10000 + 1). It matches the default settings of Seurat’s NormalizeData() function, aside from differences in floating-point error. This method is not recommended because its fixed target of 10,000 makes the implicit pseudocount depend on the dataset’s sequencing depth^22^, and because it performs worse in the benchmarks of the papers discussed below.

The next-simplest approach, (“log1pPF”), is the default. Instead of rescaling every cell to the same total of 10 thousand, it rescales each cell to the mean library size across all cells. (Library size refers to the sum of a cell’s counts across all genes.) This approach of dividing each cell’s counts by its library size relative to the mean is sometimes called “proportional fitting” (PF)^23^, and has been recommended^22^ as an improvement over rescaling to a fixed total, like logCP10k normalization does. Scanpy’s normalize_total() function uses this method with a slight variation: it uses median instead of mean to define the relative library size.

The most complex approach (“PFlog1pPF”) takes the output of log1pPF normalization and applies an additional round of proportional fitting after the log-transformation. This approach has been recommended^23^ based on the argument that log1pPF normalization does not fully normalize for read depth, because the log transform partially undoes the first round of proportional fitting.

Normalization is typically the memory bottleneck of a brisc workflow, because the normalized and unnormalized data must briefly coexist in memory. Fortunately, the sparse matrix indices, which are typically twice as large as the counts themselves (since they must be stored as 64-bit integers for large datasets), do not change, so there is only a ∼33% spike in memory usage at this step. Even so, brisc supports in-place normalization in the common scenario where the raw counts are stored as 32-bit floats, to avoid this bottleneck.

### Principal component analysis (PCA)

brisc computes the top principal components (50 by default) of the highly-variable genes, via a parallel implementation of the Implicitly Restarted Lanczos Bidiagonalization Algorithm (IRLBA), the same approximate singular vector decomposition algorithm used by Seurat’s RunPCA(). brisc runs PCA directly on the sparse version of this matrix. It standardizes each gene to zero mean and unit variance implicitly, by folding the scaling into IRLBA’s matrix-vector products. (The same implicit standardization is implemented in the irlba R package used by Seurat’s RunPCA(), although Seurat does not make use of it.) brisc uses custom matrix-vector kernels that perform reductions over fixed block sizes for deterministic results across thread counts.

Standard approximate PCA algorithms are difficult to parallelize because they spend almost all their time on matrix-vector (or matrix-matrix) products, half of which involve the full input matrix and half of which involve its transpose. Sparse matrices come in two main flavors, row-major or compressed sparse row (CSR) format and column-major or compressed sparse column (CSC) format. Transposing a CSR matrix yields a CSC matrix, and vice versa. Sparse matrix-vector products parallelize cleanly in CSR format but not CSC, so with a single copy of the matrix half of the products are always stuck single-threaded, and become the overwhelming bottleneck of the entire algorithm.

brisc’s innovation is to create both a CSR and a CSC copy of the input matrix. Half of the matrix-vector products use the CSR, and half use the CSC, so both can run in parallel. The runtime cost for creating the copy is dwarfed by the time savings during PCA. Because only highly variable genes are copied, the memory cost is also acceptable, staying below the memory usage of normalization across all datasets tested. Single-threaded runs skip the copy by default, which is about 2× faster, but match_parallel=True can be specified to force single-threaded runs to use the dual-matrix implementation and give identical numerical results to the parallel version.

PCA can be run jointly across several datasets in one call. brisc computes one common set of components based on their shared highly variable genes and returns each dataset with its own scores; no manual merge and split is required.

### Nearest neighbors

brisc finds each cell’s nearest neighbors (20 by default) in principal component space. It uses an approximate search based on the inverted file index (IVF) strategy implemented in the Facebook AI Similarity Search (FAISS) library^13^. The output is a compact dense array of neighbors and their squared distances, which is more memory-efficient than the sparse matrices Scanpy and Seurat return.

IVF-based search operates in two steps. First, all cells are clustered with k-means clustering, using a large number of clusters – the default number for large datasets (of over 640,000 cells) is four times the square root of the number of cells, rounding up. This step is based on scikit-learn’s implementation of the standard Lloyd’s k-means algorithm. Second, each cell’s nearest 64 clusters (by default) are exhaustively searched for neighbors using brute-force Euclidean distance calculations.

IVF-based search is highly accurate – brisc misses a true neighbor only when it falls outside the cell’s 64 nearest clusters – and parallelizes well. Despite being brute-force, because it scans only a small fraction of cells, it stays far faster than a full exhaustive search. brisc also implements three interlinked optimizations which collectively speed up the search by about an order of magnitude over FAISS’s IVF implementation, without changing the result.

First, brisc sorts cells by cluster and orders the clusters along their first principal component, so that consecutive cells are nearby in principal component space. The sorted order implicitly encodes the inverted file index (an index mapping each cluster to the list of cells it contains), avoiding the need to build one explicitly.

Second, instead of processing each cell one at a time, brisc groups consecutive cells into blocks and processes them together. Because consecutive cells are nearby in principal component space (the first optimization), they will tend to share many of their 64 nearest clusters, so the union of clusters the block needs to search is sparse – far smaller than the theoretical worst-case scenario of 64 × block size. brisc exploits this property by gathering that union, then searching each cluster in the union against all block members that have it among their 64 nearest clusters. This amortizes the cost of streaming the cluster’s cells from memory across all the cells in the block that need them.

Third, brisc exploits the fact that the Euclidean distance between a query cell Q and a candidate cell C can be rewritten using the identity ||Q - C||² = ||Q||² - 2Q·C + ||C||², where ||Q||² and ||C||² are the squared L2 norms of Q and C and Q·C is a matrix multiplication. This allows the squared norm ||Q||² to be ignored, since it is the same for every candidate and does not affect how the candidates rank against each other. More importantly, because cells are grouped into blocks (the second optimization), the Q·C term can be computed via BLAS matrix multiplication, which is highly optimized, using cache-blocking and tuned microkernels that a pairwise distance loop cannot match. The same identity also accelerates the computation of distances from cells to centroids, both in the nearest-neighbors search and in the k-means clustering.

brisc also implements a fourth, smaller optimization: once the distances have been calculated in this blockwise fashion, it is necessary to scan across them to find which candidate centroids or cells are closest. brisc scans cell candidates in groups of 64 and finds the smallest distance in each group, which processors can vectorize using single instruction multiple data (SIMD) instructions. If even the nearest candidate does not beat the current worst neighbor, brisc skips the entire group.

All steps in brisc that use BLAS, including this one, use single-threaded BLAS, since multithreaded BLAS is often nondeterministic and is generally slower than keeping BLAS single-threaded and instead parallelizing the loop that calls it.

### Shared nearest neighbors

Prior to Leiden clustering, brisc uses the nearest neighbors to calculate a shared-nearest-neighbor (SNN) graph. Edges are weighted by Jaccard index (the number of neighbors two cells share, divided by the total number of distinct neighbors between them), matching Seurat’s definition. Scanpy instead derives connectivity from the UMAP manifold. Weak edges are pruned: two cells are linked only if they share at least 3 neighbors (matching Seurat’s prune.SNN = 1/15 when there are 20 neighbors).

brisc achieves this using an optimized parallel approach that outperforms Seurat’s implementation in both speed and memory efficiency. To find shared neighbors, Seurat performs a sparse matrix multiplication (SNN · SNN^T^), which forces it to instantiate the entire unpruned graph in memory before filtering out weak edges. In contrast, brisc uses a two-pass approach. By traversing reverse-neighbor lists and sorting shared candidates in thread-local, heavily-reused buffers, brisc calculates the exact number of pruned edges in the first pass. This allows it to allocate the final sparse matrix of shared neighbors to its exact size before filling it in the second pass, avoiding large intermediate memory spikes.

### Clustering

brisc clusters cells by running Leiden clustering^24^ on the SNN graph. It optimizes the same Reichardt-Bornholdt Potts objective^25^ as Seurat and Scanpy; at the default resolution of 1, this is equivalent to optimizing modularity. Leiden operates in three iterative phases: a local move phase that greedily reassigns nodes to neighboring communities to maximize the objective, a refinement phase that partitions these communities into “refined” communities to guarantee well-connected components, and an aggregation phase that collapses the refined communities into a supergraph. This cycle repeats on the newly aggregated supergraph until no further merges can be made.

As a graph algorithm where nodes depend on the current state of their neighbors, which may be modified by other threads mid-computation, Leiden clustering is inherently difficult to parallelize deterministically, and we are not aware of any deterministic parallel implementations. (Our implementation is loosely based on GVE-Leiden^26^, an efficient but non-deterministic parallel implementation.) As a result, brisc leaves the Leiden clustering itself single-threaded, and instead parallelizes across resolutions. Since it is often recommended to run Leiden clustering at a variety of resolutions (e.g. 5 to 10) to explore clusters of different granularities, this approach still achieves meaningful parallelism without sacrificing reproducibility.

Even single-threaded, brisc’s Leiden clustering is several times faster than scanpy’s igraph^14^ backend. (igraph is Scanpy’s upcoming default backend, and is itself faster than the current leidenalg default.) The difference is primarily due to systems engineering, across four areas: memory layout, allocation strategy, prefetching, and arithmetical optimizations.

**Memory layout:** igraph initializes and traverses a dynamic incidence list, which requires an extra pass to build and scatters edges non-contiguously in memory. brisc bypasses this by operating directly on the native CSR arrays (data, indices, indptr) of the SNN graph, maximizing spatial locality.

During aggregation, igraph collapses communities by continuously pushing elements into dynamically resizing vectors, which is slow and fragments memory. brisc instead uses a counting sort: it first counts how many nodes belong to each community to compute prefix offsets, then in a single pass places every node directly into a pre-allocated flat array so that nodes in the same community are contiguous. Once grouped, a linear scan over these contiguous blocks rapidly sums inter-community edge weights to construct the new supergraph.

**Allocation strategy:** igraph dynamically creates a new graph object on every pass of the aggregation phase and frees the old one. brisc instead allocates each buffer once: the per-node buffers before the first iteration, and the two sets of supergraph buffers at the end of the first two iterations, once their maximum sizes are known. Because the number of nodes, edges and communities shrinks on each iteration, later iterations operate on increasingly smaller fractions of these buffers. For the supergraph itself, brisc uses a ping-pong buffer strategy: one set of buffers (data, indices, indptr) is used on even-numbered iterations, a second on odd-numbered iterations. This lets the algorithm safely read the current graph’s structure while simultaneously writing the new supergraph, eliminating memory copies or dynamic allocations inside the solver loop.

brisc further reduces its memory footprint by sharing buffers across mutually exclusive phases: for example, the buffer allocated for the circular queue (used to track which nodes need to be revisited) during the local move phase is reused during refinement to map community labels to contiguous indices, again during aggregation to track which neighboring communities have been evaluated, and once more at the end to relabel final clusters.

**Prefetching:** Graph traversal is heavily memory-bound due to unpredictable pointer chasing – looking up a node, then its neighbor indices, then those neighbors’ community labels – which is a pattern hardware prefetchers cannot predict. During the local move phase, brisc issues explicit, staggered prefetch hints (builtin_prefetch() on Linux and Mac) down this chain: row pointers 24 nodes ahead, edge indices 16 nodes ahead, and node state (community assignment, weighted degree, community weight) 8 nodes ahead. The refinement phase visits nodes in index order, which the hardware prefetcher handles on its own, and the aggregation phase prefetches node state 8 nodes ahead. During neighbor traversal in all three phases, brisc additionally prefetches community labels of adjacent neighbors 16 edges ahead. This multi-tiered prefetching effectively hides memory latency and yields a ∼20% speedup on large datasets.

**Arithmetic:** During the local move and refinement phases, each node evaluates whether to join a neighboring community. brisc hoists an invariant term – the resolution parameter scaled by the node’s weighted degree outside the loop over neighbors – computing it once per node instead of once per neighbor. In the local move phase, it also hoists a second invariant term, the baseline objective (the cost of the node staying in its current community), and initializes the best observed objective to this baseline rather than zero, which eliminates a subtraction from every iteration of the innermost loop. igraph recomputes these invariant terms for every neighbor examined, resulting in millions of redundant floating-point operations on large datasets.

Finally, brisc rethinks how the accumulator is reset between nodes. To identify which unique communities a node connects to, igraph maintains a separate tracking array and checks a bitset on every edge to prevent duplicates – introducing branch mispredictions into the hottest loop of the algorithm. brisc drops the bitset and tracking array entirely during accumulation, summing edge weights without any branching. To evaluate and reset, it simply loops over the node’s edges a second time: if a community’s accumulated weight is nonzero, brisc evaluates the objective and resets the entry in place. This trades a few cheap sequential memory reads for the elimination of costly branch mispredictions and bitset logic.

As a quality-of-life postprocessing step, brisc merges cells in “tiny” clusters (with fewer than 5 cells, by default) into the community of their nearest neighbor that is not in a tiny cluster.

### Embedding

brisc supports three methods for computing two-dimensional embeddings for visualization. The default in both Scanpy and Seurat, Uniform Manifold Approximation and Projection (UMAP), is the established standard^9^. It preserves local structure well but tends to distort the global arrangement of clusters.

Pairwise Controlled Manifold Approximation (PaCMAP)^7^ is a newer method that preserves both local and global structure simultaneously. It positions cells by balancing attractive and repulsive forces across three types of cell pairs: nearby neighbors are pulled together to capture local detail, mid-distance pairs provide a global scaffold, and distant pairs are pushed apart.

Pairwise Controlled Manifold Approximation with Local Adjusted Graph (LocalMAP) is a variant of PaCMAP that periodically re-samples its repulsive pairs from cells that have drifted close together during optimization^8^. This aims to prevent distinct clusters from collapsing into one another in the final layout, a failure mode that can obscure biologically meaningful structure.

PaCMAP and LocalMAP are substantially faster than UMAP (with PaCMAP being the fastest) and support deterministic parallel execution; UMAP does not. To achieve deterministic parallelism in PaCMAP and LocalMAP, brisc introduces a structural optimization not found in the original implementations of these algorithms. Traditionally, these algorithms compute gradients by iterating over defined pairs of cells (i, j) and symmetrically updating the gradients for both cells simultaneously. In a multithreaded context, this creates race conditions: multiple threads processing different pairs might attempt to update cell j’s memory at the exact same time, which forces a choice between using computationally expensive memory locks or accepting non-deterministic or thread-varying outputs.

To resolve this, brisc transforms the 2D array of sampled pairs into a symmetric sparse adjacency list (similar to a Compressed Sparse Row matrix) prior to optimization. Every (i, j) relationship is recorded twice – once for i and once for j – so that each cell maintains a complete, independent array of its interacting partners. During the gradient update step, execution is parallelized across cells rather than pairs. A thread assigned to cell i iterates over all of i’s interacting partners to accumulate the attractive and repulsive forces, but it applies the resulting gradient update only to cell i. Because no thread ever modifies the gradient of a cell assigned to another thread, memory writes are completely isolated. This design allows brisc to execute lock-free, highly parallelized gradient descent while guaranteeing perfectly reproducible, deterministic results regardless of thread execution order. However, since it has some computational overhead, it is turned off for single-threaded execution unless match_parallel=True is specified, like for PCA.

Finally, brisc also provides a parallel UMAP implementation using the Hogwild! algorithm for lock-free asynchronous gradient descent^15^, enabled by setting hogwild=True. This is the only non-deterministic step in brisc. brisc’s UMAP implementation closely follows umap-learn’s, but uses a rejection sampling-style approach rather than deterministic scheduling to decide which edges to visit on each epoch: each edge is visited with probability equal to its weight. The two approaches give each edge the same expected number of updates, but brisc’s approach does not need to store scheduling metadata for every edge, which simplifies the code and reduces memory traffic. brisc also uses explicit prefetching (a technique described in the Clustering section, above) for both positive neighbors and negative samples.

### Marker genes

brisc implements marker-gene finding to identify the genes that distinguish each cell type from all the others. Conventional tools find markers via differential-expression tests that compare expression distributions between groups (like Scanpy’s rank_genes_groups() and Seurat’s FindAllMarkers()). brisc instead scores each gene by two simpler quantities: its detection rate (the fraction of cells with non-zero counts for the gene) in the cell type, and the fold change in that detection rate between the cell type and all other cell types pooled together. This adapts Fischer and Gillis (2021)^16^, who rank markers by AUROC and fold change in detection rate. brisc substitutes detection rate for AUROC because it is far cheaper to compute and, by the authors’ own results, detection rate and AUROC are almost perfectly correlated.

The two metrics brisc uses to select markers (detection rate and fold change in detection rate) trade off against each other – a gene expressed in nearly every cell of a type (high detection rate) is usually expressed in other types too (low fold change), and vice versa – so no single gene is best. brisc selects every gene on the Pareto front of the two metrics: the genes that no other gene beats on both detection rate and fold change at once. Candidate markers must also clear minimum thresholds on both metrics: 25% detection rate and 2-fold change by default.

This approach is several orders of magnitude faster than conventional tools, partly due to its inherent simplicity and partly due to a number of implementation choices. First, detection rates are counts of non-zero entries per cell-type group computed directly from the indices of the sparse matrix, similar to pseudobulking.

Second, brisc avoids recomputing the background group for the fold-change calculation (a gene’s detection rate in all other cell types pooled together) from scratch for each cell type. Instead, it calculates each gene’s detection rate in all cell types (which is just the total number of non-zero entries per column of the sparse matrix) once, then merely subtracts off the foreground (the gene’s detection rate in the cell type of interest) for each cell type to obtain the background.

Third, computing the Pareto front is naively O(N^2^) in the number of genes; brisc speeds this up by first discarding any gene that fails the minimum thresholds, then parallelizing across cell types, so that each thread computes the Pareto front of one cell type independently and threads never need to coordinate. Within a cell type, each candidate gene is compared against every other gene until one is found that matches or beats it on both metrics, and genes that tie on both metrics are resolved deterministically so that only one of them is kept.

Finally, for CSR-format count matrices, when the number of available CPU threads exceeds the number of cell types, parallelizing over cell types would leave cores idle. brisc instead parallelizes over cells in chunks, accumulates counts into thread-local scratch buffers, and combines them with a parallel reduction.

### Harmonization

brisc implements the Harmony2 algorithm^17^ to remove batch effects across datasets or batches. Harmony2 works by iterating two steps: a soft clustering step that groups cells by transcriptional similarity while penalizing clusters dominated by a single batch, and a ridge regression step that estimates and removes each batch’s contribution to the PC coordinates within each cluster, which is computed analytically via an arrowhead matrix inverse approach. However, brisc departs from the reference R implementation of Harmony2 in three major respects.

First, brisc supports only a single batch covariate: when both datasets and batches within them are specified, each combination is treated as a distinct batch. This restriction lets it represent batch assignments as a 1D vector of integers rather than a one-hot encoded design matrix. The reference R implementation uses a sparse matrix encoding of batch assignments, which it multiplies against cell embeddings throughout the algorithm. brisc discards this abstraction entirely: batch membership is stored as a single integer per cell, and that integer is used as a direct array index throughout the Cython inner loops. This batch vector is used pervasively throughout the algorithm, leading to contiguous, cache-friendly memory accesses in place of sparse matrix operations, as well as a reduced memory footprint. The restriction also guarantees that the ridge regression has the arrowhead structure that the reference R implementation exploits when there is a single covariate, so each cluster’s regression has a closed-form solution based on summing the cluster probability-weighted PCs within each batch. Whereas the reference R implementation solves the regression separately for each cluster, making several passes over all datasets’ Harmony embeddings for each cluster, brisc groups cells by batch and computes these sums for every batch and cluster at once with dense matrix multiplications, making only two passes in total (one to compute the sums and one to apply the correction) regardless of the number of clusters.

Second, brisc introduces parallelism into an inherently sequential algorithm via a nested “chunks-within-blocks” strategy. The reference R implementation processes cells one block at a time (5% of the data per block by default), updating the observed and expected cluster-by-batch co-occurrence matrices (O and E) after each block. Because each block’s update depends on the result of the previous block, no parallelism is possible at the block level without changing the algorithm. brisc preserves the sequential block structure but subdivides each block into fixed-size chunks of cells (by default, 256 cells or the number of batches rounded up to a multiple of 16, whichever is larger), which are processed in parallel. Each thread computes its chunks’ contributions to O and E independently; the contributions are then summed and applied to the global matrices once per block. Each chunk’s assignments are updated against O and E with only that chunk’s own contribution removed, whereas the reference R implementation removes the whole block’s. The ridge regression step, which has no sequential dependencies, instead uses a separate partition of the cells into “slabs” of consecutive cells, independent of the blocks and chunks. Each slab contains roughly max(chunk size, number of batches × number of clusters) cells, which lets it store its partial sums for the regression in its own rows of the embedding matrix rather than in additional memory. Slabs are processed in parallel and NUMA-locally. Because chunk and slab boundaries are fixed and independent of thread count, results are numerically identical regardless of how many threads are used.

Third, brisc avoids separate passes over the data that the reference R implementation makes. Both implementations alternate between refining cluster assignments and updating the Harmony embeddings, running multiple cluster refinement iterations per embedding update step. The reference R implementation stops cluster refinement early once the average value of the objective function (a sum of a k-means error term, an entropy term, and a batch-diversity penalty) over the three most recent refinement iterations converges. However, it only starts checking whether to early-stop on the fifth iteration, so at the default of 4 iterations the check never runs. It nonetheless calculates the objective function on every iteration. By default, brisc instead skips the early-stopping check and always runs exactly 4 refinement iterations. This lets it compute the objective function only on the final iteration, where it is used to decide whether the harmonization as a whole is finished, and as part of the same pass that updates the assignment probabilities rather than in a separate pass. Similarly, brisc recomputes the assignment probabilities from the updated embeddings during the same pass that updates the embeddings, rather than in a separate pass.

brisc also differs from the reference R implementation in minor ways unrelated to its parallel structure. To reduce runtime, it initializes k-means with random rather than k-means++ seeding, adds early stopping to the k-means, and shuffles cells once at the start rather than on every pass. It also retains the original Harmony algorithm’s^27^ behavior of recomputing each cluster’s centroid after every refinement iteration, rather than holding the centroids fixed and resetting them to the regression intercepts after each PC correction step.

### Label transfer

brisc implements label-transfer to carry cell-type labels from a labeled reference dataset onto an unlabeled query, once both have been placed in a shared space by brisc’s harmonization. For each query cell, it finds the nearest reference cells in the Harmony embedding – using the same fast IVF-based search described in the nearest-neighbors section, but pointed at the reference dataset rather than the query dataset itself – and assigns the most common label among them (20 neighbors by default), reporting the fraction that agree as a confidence. Ties are broken in favor of the globally most common cell type in the reference. The search and vote are parallelized, and there is an option to return each cell’s runner-up label as well.

### Pseudobulking

Pseudobulking collapses a single-cell dataset into bulk-like profiles by summing the raw counts of all cells that share a sample and cell type. The result is a Pseudobulk dataset containing one count matrix per cell type, with samples as rows and genes as columns. Counts are summed as integers, so totals are exact with no floating-point error. The summation operates directly on the sparse matrix, in parallel across sample-by-cell-type groups (for CSR count matrices) or genes (for CSC), into a preallocated array – aggregating even very large datasets in a single pass without densifying or copying.

brisc also infers sample-level metadata automatically. For each sample, brisc identifies the cell-level metadata columns that are constant across all of that sample’s cells (e.g. donor age, sex, condition) and carries these into the output Pseudobulk dataset, discarding columns that vary from cell to cell. brisc also automatically injects a column into the Pseudobulk dataset’s metadata indicating the number of cells that went into each pseudobulk profile for a given sample and cell type.

### Differential expression

brisc implements a highly flexible pseudobulk differential expression testing framework based on limma-voom^28,29^, which it interfaces with via the ryp Python-R bridge. The design is specified as an R formula (e.g. ∼ disease_status + age + sex) which pulls the relevant columns from the Pseudobulk dataset’s metadata. brisc recommends adding log2(num_cells) + log2(library_size) to the formula to avoid confounding due to differences in number of cells or library size across samples.

brisc implements the standard limma-voom pipeline: library sizes are calculated via trimmed mean of M-values (TMM) normalization^30^, the design matrix is built with R’s model.matrix(), mean-variance precision weights are estimated with voom, these weights are used to fit a weighted linear model with limma’s lmFit(), and per-gene variances are moderated with empirical Bayes (eBayes()). Contrasts can be specified, and are handled with limma’s makeContrasts() and contrasts.fit(). brisc uses voomByGroup^18^ rather than plain voom whenever the comparison is categorical, such as when comparing cases versus controls or providing contrasts. Cases and controls can have different mean-variance trends, so fitting one global trend misweights genes; voomByGroup fits a separate trend per group for better-calibrated weights.

All of the above is exposed through a single function call. High-level wrappers for complex pipelines tend to fail in one of two ways: they abstract away decisions the user needs control over, or they expose too many options that they lack sensible defaults for. brisc aims to avoid these pitfalls, inferring reasonable defaults wherever possible while remaining fully configurable. By default, it tests the first non-intercept term in the design matrix, automatically encodes string and categorical metadata as unordered factors, and infers the appropriate sample groupings for voomByGroup directly from the formula – but every aspect of the design can be overridden per cell type. brisc also handles the inherent sparsity of single-cell data gracefully: if a cell type lacks enough samples to build a full-rank design matrix, it is skipped rather than crashing the pipeline.

### Benchmarking

#### Datasets

The three datasets are described in the Results. We used brisc to pre-convert each dataset to the same layout (an uncompressed, unchunked .h5ad file of raw counts in CSR format alongside the minimal metadata necessary for our workflows), so that every toolkit could read from the same file during benchmarking.

#### Hardware and software

All benchmarks ran on the Trillium cluster (SciNet, University of Toronto), with one benchmarking job per node, under a 24-hour time limit. The CPU node has 192 cores and 749 GB of RAM, while the GPU node has 96 cores, 752 GB of RAM and 4 NVIDIA H100 80 GB GPUs. Neither has a local disk, so datasets were read from the cluster’s networked file system.

We used brisc 0.2.3, Scanpy 1.12, Seurat 5.6.0 with BPCells 0.3.1 and rapids-singlecell 0.15.0. To compare the CPU toolkits on an equal footing, we compiled Python, R and every compiled package they use (including brisc, Seurat and BPCells) from source with the same compiler and optimization flags, linked against Intel MKL. Full software environments are given in the benchmarking code (Code availability).

#### Benchmark design

Each workflow was implemented as one script per toolkit, and each step used the toolkit’s defaults unless stated below. For an apples-to-apples comparison, we used brisc’s default quality control filters for every toolkit: at least 100 genes detected, at most 5% mitochondrial counts and non-zero MALAT1 expression. We ran each workflow three times to assess reproducibility and run-to-run variability in runtime. brisc’s multi-threaded runs used all available cores.

Seurat was benchmarked with BPCells, as recommended by Seurat’s developers for large datasets, because Seurat alone cannot load any of the three datasets. Three of Seurat’s own functions (neighbor search, clustering and pseudobulking) fail or are impractically slow at this scale, so we used BPCells’ equivalents instead. Otherwise Seurat and BPCells ran at their defaults, under which BPCells’ streaming operations are single-threaded. Scanpy also ran at its defaults, except that Leiden used the igraph backend (Scanpy’s announced future default). rapids-singlecell ran on the GPU node’s four GPUs according to its multi-GPU tutorial, with the dataset read in chunks into a Dask array distributed across the GPUs.

#### Workflows

The standard workflow (Fig. 1b) selected 2,000 highly variable genes and 50 principal components, clustered at five Leiden resolutions (0.25 to 2), and (for an apples-to-apples comparison) used the cell-type calls that came with the dataset for marker-gene finding, rather than the ones found by each toolkit’s Leiden clustering. The toolkits differ at several steps. brisc and Scanpy select variable genes within each donor (or each mouse in PanSci), whereas Seurat and rapids-singlecell do not (Seurat lacks this functionality and rapids-singlecell’s implementation of it is extremely slow). On the GPU node, brisc also ran without batches to match rapids-singlecell. brisc uses LocalMAP for embedding while the other toolkits use UMAP, although we also report brisc’s PaCMAP and UMAP results (Extended Data Fig. 6). Markers were found by brisc’s Pareto-front method, Scanpy’s t-test, Seurat’s Wilcoxon test (through presto) and rapids-singlecell’s logistic regression.

For label transfer, we split each dataset by condition into an annotated reference and a query: SEA-AD’s neurotypical reference donors versus its remaining donors, PBS-treated versus cytokine-treated cells in Parse, and wild-type versus immunodeficient mice in PanSci. brisc harmonizes the reference and query before transferring labels from each query cell’s 20 nearest reference cells, Scanpy transfers them with ingest and Seurat with transfer anchors.

We tested differential expression between Alzheimer’s disease and control donors in SEA-AD, between interferon-gamma-treated and PBS-treated cells in Parse, and between 23-month-old and 6-month-old wild-type mice in PanSci, with cells outside each comparison excluded at the QC stage in every toolkit. Accordingly, we pseudobulked across donors in SEA-AD, donor-treatment combinations in Parse, and mouse-organ combinations in PanSci. Differential expression models adjusted for APOE ε4, sex, age and post-mortem interval in SEA-AD, for donor in Parse and for sex in PanSci, and in every dataset for number of cells and library size. For Scanpy, which supports pseudobulking but does not have functionality for QC or differential expression on pseudobulks, we used the decoupler package for QC and pydeseq2 for differential expression, with the same covariates as brisc. For Seurat, we ran DESeq2 through Seurat’s own interface, which supports neither covariates nor pseudobulk-level QC.

#### Memory benchmarking

Peak memory usage was measured by a monitor process that sampled the toolkit’s resident set size every 50 ms, with a step’s peak memory usage being defined as the largest sample taken while it ran. brisc’s parallel loading required a different measure, because its worker processes read the file into memory shared with the main process, so a sum of resident set sizes would count shared pages several times. During loading, each sample was therefore the proportional set size summed over all of brisc’s processes, a strategy which counts each page exactly once. Memory is reported in binary units (1 GB = 2^30^ bytes).

#### Accuracy and reproducibility

Nearest-neighbor recall was measured against exact neighbors computed by brute force from each toolkit’s own principal components (Fig. 5). The agreement between each toolkit’s Leiden clustering results and the cell-type calls that came with each dataset was quantified via the balanced adjusted Rand index^31^, which weights every cell type equally regardless of the number of cells it contains. Clusters of fewer than 100 cells were not counted, because Seurat’s pruning of the shared-nearest-neighbor graph leaves it with many singleton clusters.

Our reproducibility analysis compared the three runs pairwise, with the worst difference between any pair shown in Fig. 6. Each step used a different reproducibility metric. When comparing which cells were retained by QC, we looked at overlap according to the Jaccard index. When comparing which genes were selected as highly variable, we looked at how many of the top 2000 genes were selected by both runs. For principal components, we quantified the root-mean-square difference of the worst component (after sign alignment) relative to its standard deviation. For nearest neighbors, we looked at what fraction of each cell’s neighbors were the same between runs, on average.

Because clustering is unsupervised, the same groups of cells may be assigned different cluster labels in different runs. Clusters therefore cannot be compared by their assigned labels (e.g., cluster 3 in one run is not necessarily cluster 3 in another). Instead, for each pair of runs, we matched clusters using the Hungarian algorithm^32^ to find the optimal one-to-one correspondence between clusters in one run and clusters in the other. We then quantified reproducibility using cluster agreement, defined as the fraction of cells whose cluster assignments agreed between the two runs after this matching, together with the adjusted Rand index and the number of clusters.

## Data availability

All three benchmarked datasets are publicly available. The SEA-AD middle temporal gyrus single-nucleus RNA-seq data (SEAAD_MTG_RNAseq_final-nuclei.2024-02-13.h5ad) and donor metadata were downloaded from the Allen Institute’s SEA-AD data repository (https://sea-ad-single-cell-profiling.s3.amazonaws.com/index.html; https://portal.brain-map.org/explore/seattle-alzheimers-disease). The Parse Biosciences 10 million PBMC cytokine dataset (Parse_10M_PBMC_cytokines.h5ad) was downloaded from https://www.parsebiosciences.com/datasets/10-million-human-pbmcs-in-a-single-experiment/. The PanSci mouse atlas was downloaded from the Gene Expression Omnibus under accession GSE247719.

## Code availability

brisc is available at https://github.com/briscverse/brisc. Documentation is available at https://brisc.run. Benchmarking scripts are available at https://github.com/keon-arbabi/brisc-benchmarking.

## Acknowledgements

We thank Shreejoy Tripathy and the members of his group for testing and providing feedback on an early version of brisc.

## Author contributions

KA and MW conceived the study. MW implemented brisc with help from KA, TL, JJ and HH. KA wrote and ran the benchmarks, with help from ML. KA and MW designed the figures and wrote the manuscript, with feedback from TL, GDB and VM. MW supervised the work. All authors read and approved the final manuscript.

## Competing interests

The authors declare no competing interests.

## Extended Data Figures

**Extended Data Fig. 1.**
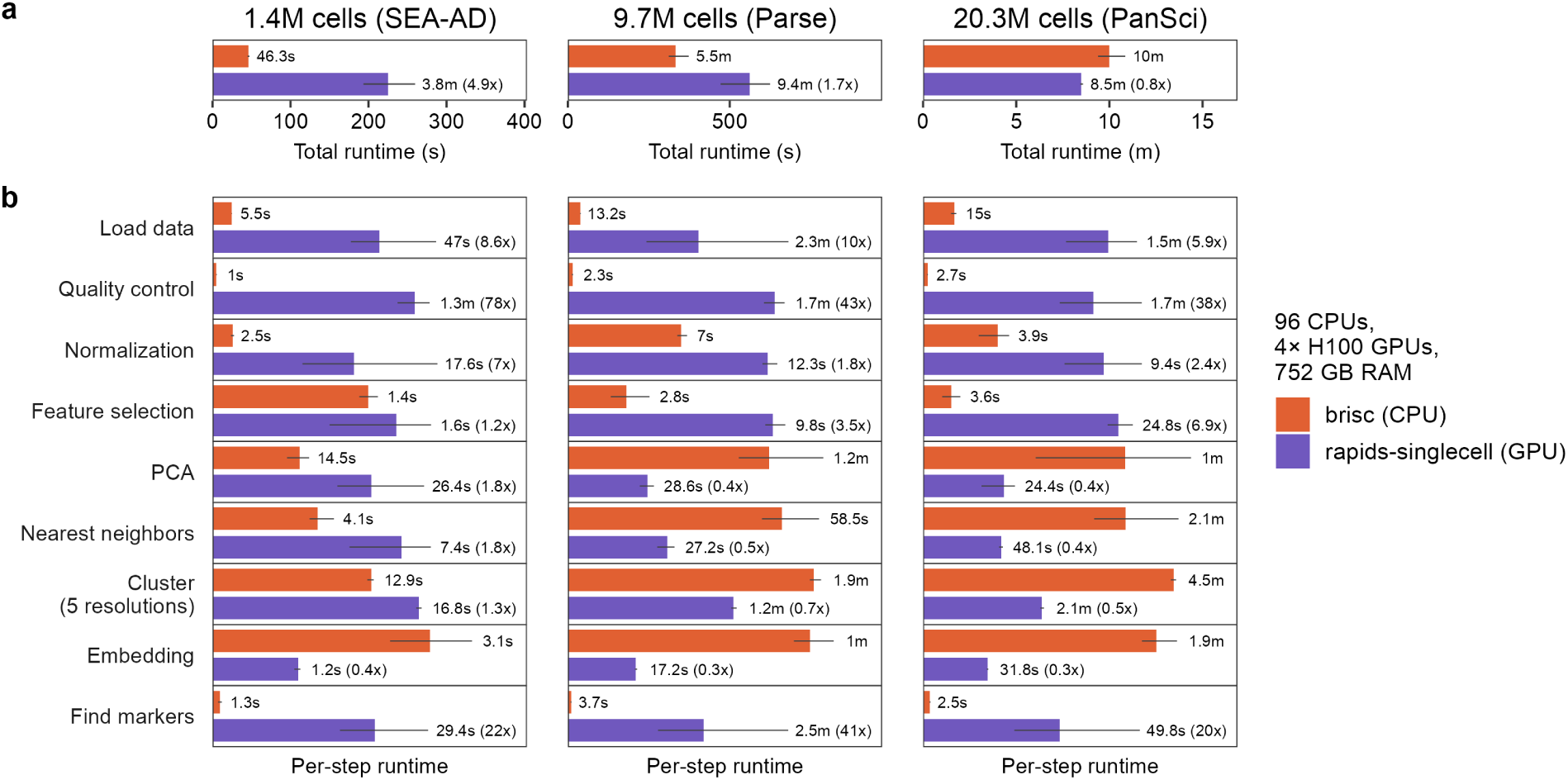
Runtime of the standard workflow on CPUs versus GPUs. **a**, Total runtime of the standard workflow on three datasets (columns), on a GPU node (96 cores, 4 NVIDIA H100 GPUs) where brisc used the CPUs and rapids-singlecell used the GPUs. **b**, Runtime of each step of the standard workflow (rows), with runtime axes scaled independently for each step. Bars show the mean runtime of three runs; error bars show the range. Each bar is labeled with its runtime, and rapids-singlecell’s bars also include its runtime as a multiple of brisc’s, so values above 1 indicate that brisc was faster. brisc computes the embedding with LocalMAP and rapids-singlecell with UMAP.

**Extended Data Fig. 2.**
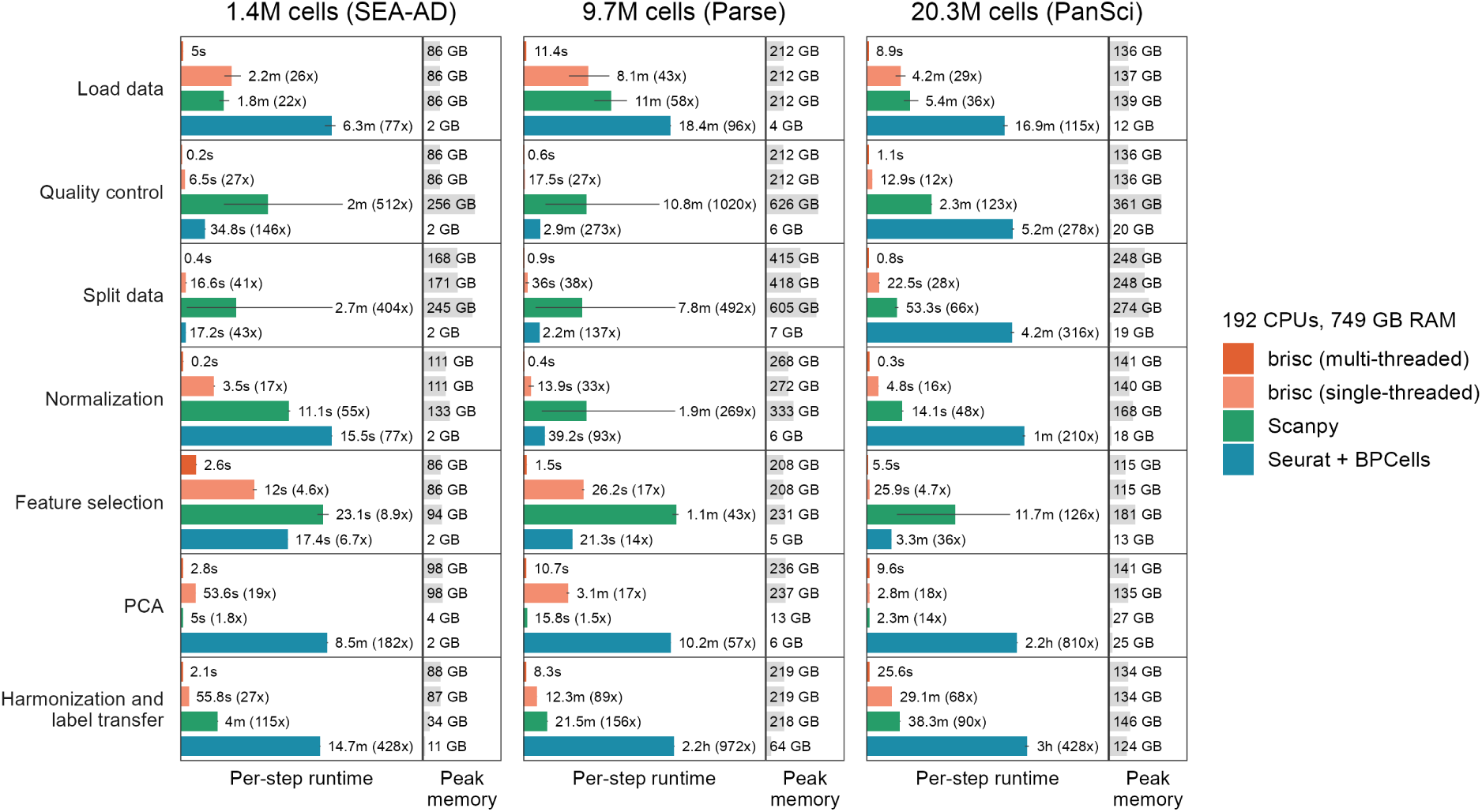
Per-step runtime and peak memory usage of the label-transfer workflow. Runtime and peak memory of each step of the label-transfer workflow (rows) on three datasets (columns), from the same runs as in Fig. 3. Display conventions as in Fig. 4. Step implementations differ between toolkits. brisc computes HVGs and PCA jointly for the reference and query, whereas Scanpy and Seurat compute it for the reference only. The harmonization and label transfer step is Harmony2 harmonization followed by nearest-neighbor label transfer for brisc, ingest for Scanpy and transfer anchors for Seurat.

**Extended Data Fig. 3.**
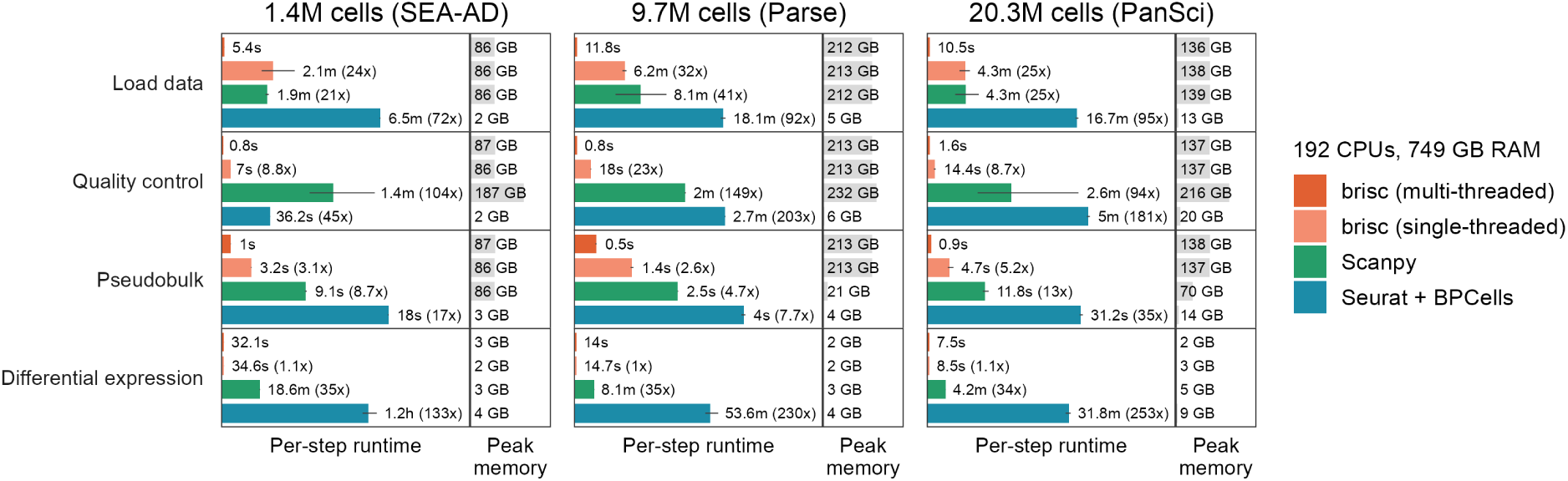
Per-step runtime and peak memory usage of the pseudobulk differential-expression workflow. Runtime and peak memory usage of each step of the pseudobulk differential-expression workflow (rows) on three datasets (columns), from the same runs as in Fig. 3. Display conventions as in Fig. 4. Step implementations differ between toolkits. The Pseudobulk step sums each sample’s counts per cell type in every toolkit, with Seurat using BPCells’ pseudobulking functionality. Quality control comprises cell-level filtering and (for brisc and Scanpy) pseudobulk-level filtering, which Seurat lacks. The Differential expression step is limma-voom with voomByGroup for brisc, pydeseq2 for Scanpy and DESeq2 for Seurat.

**Extended Data Fig. 4.**
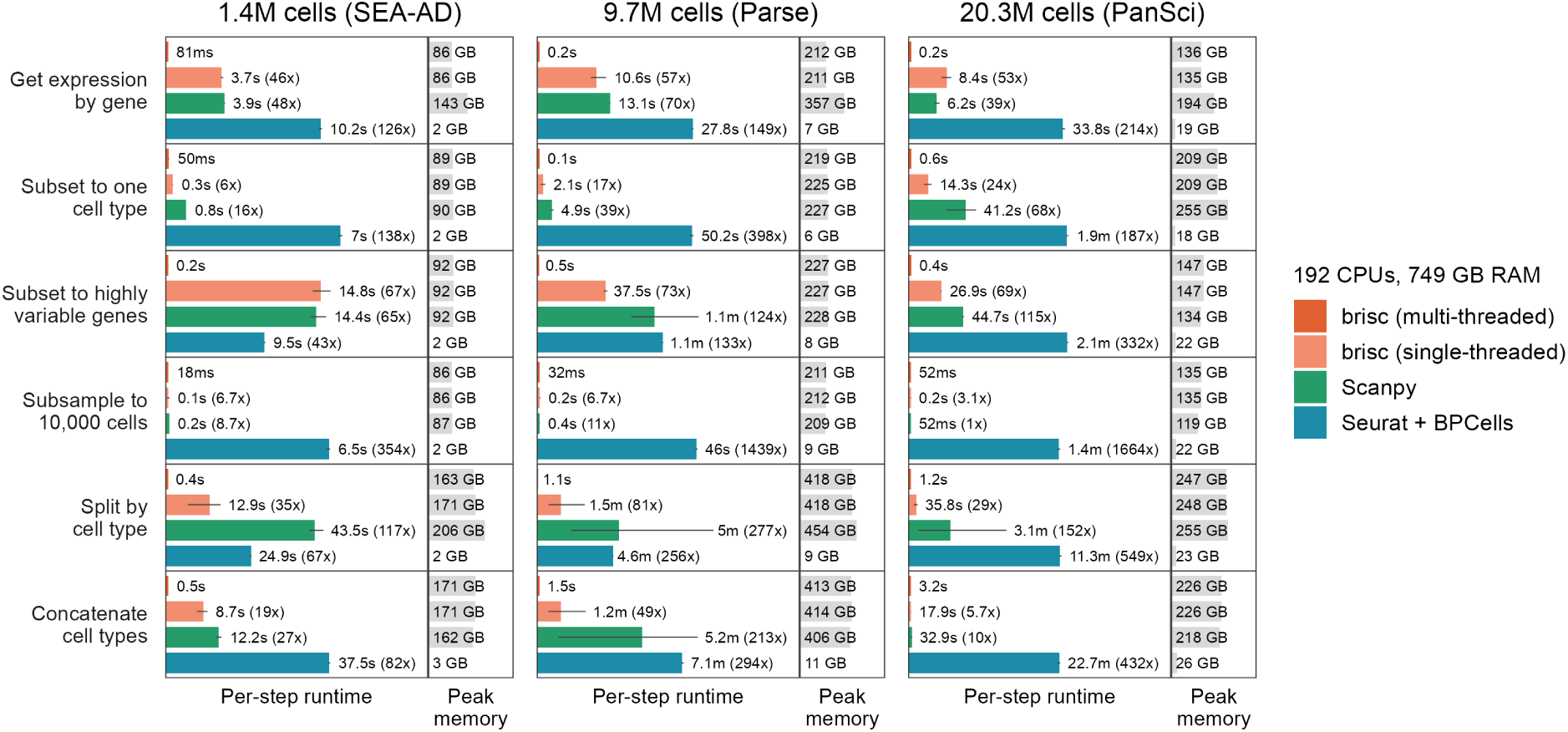
Runtime and peak memory usage of common data-manipulation operations. Runtime and peak memory of six data-manipulation operations (rows) on three datasets (columns). Display conventions as in Fig. 4.

**Extended Data Fig. 5.**
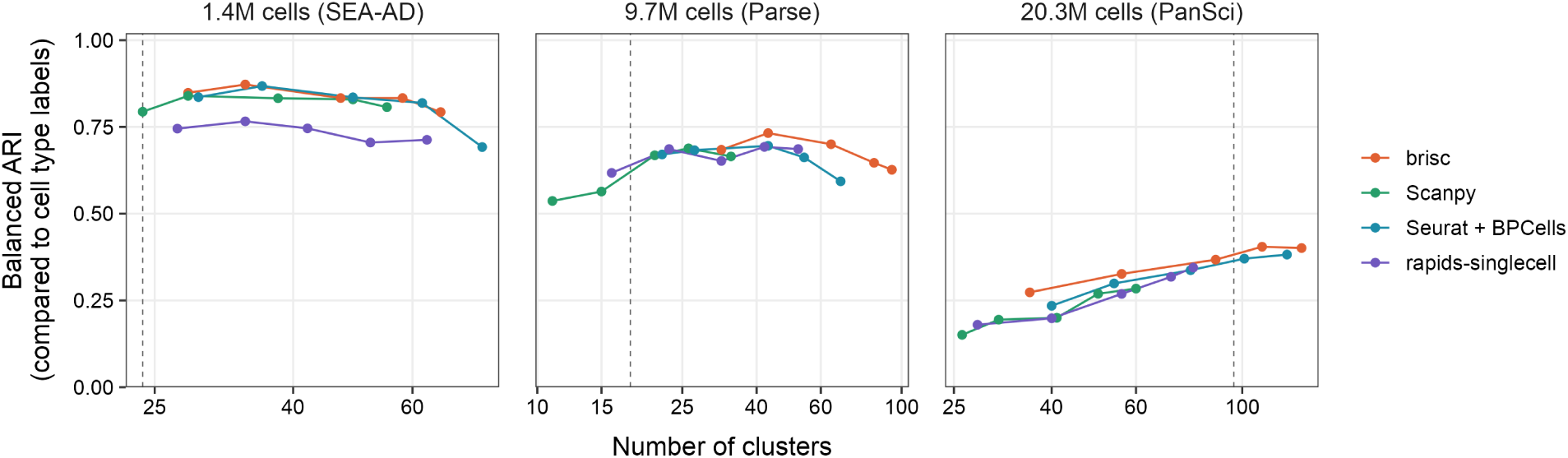
Agreement between clusters and author cell-type annotations. Balanced adjusted Rand index (ARI) between each toolkit’s Leiden clusters and the author cell-type annotations, plotted against the number of clusters at each of the five resolutions of the standard workflow (0.25, 0.5, 1, 1.5 and 2). Dashed lines mark the number of annotated cell types. The x-axes, indicating the number of clusters, are log-scaled.

**Extended Data Fig. 6.**
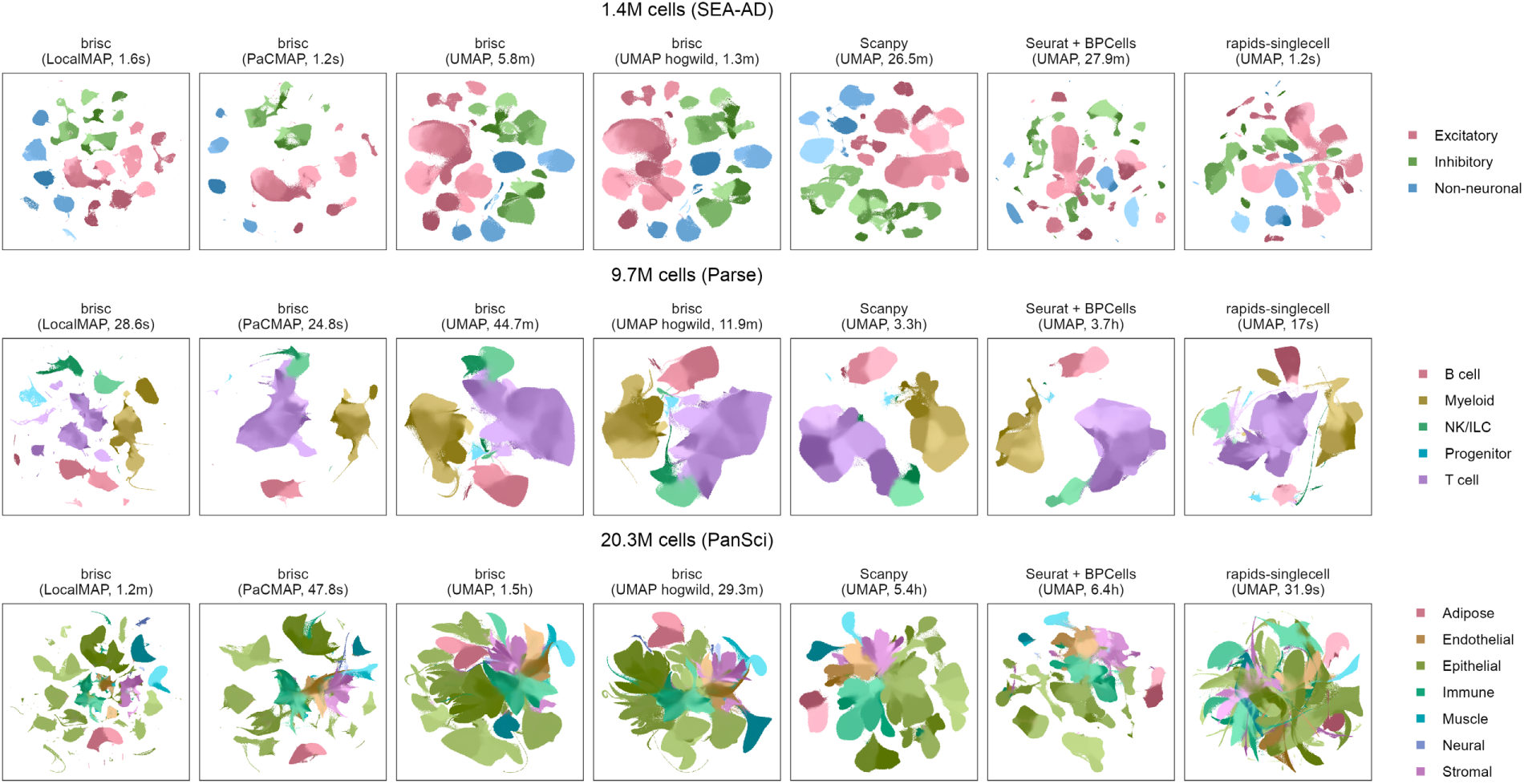
Two-dimensional embeddings across toolkits and methods. Embeddings of the three datasets (rows) computed by brisc with LocalMAP, PaCMAP, UMAP and Hogwild! UMAP (a parallel but non-deterministic variant), and by Scanpy, Seurat and rapids-singlecell with UMAP (columns), labeled with each method’s runtime in the plotted run. Cells are colored by Leiden cluster at a resolution of 1, with clusters of the same broad cell type sharing the same broad color (legend). Since each toolkit produces its own clusters, colors are not comparable across toolkits.

**Extended Data Fig. 7.**
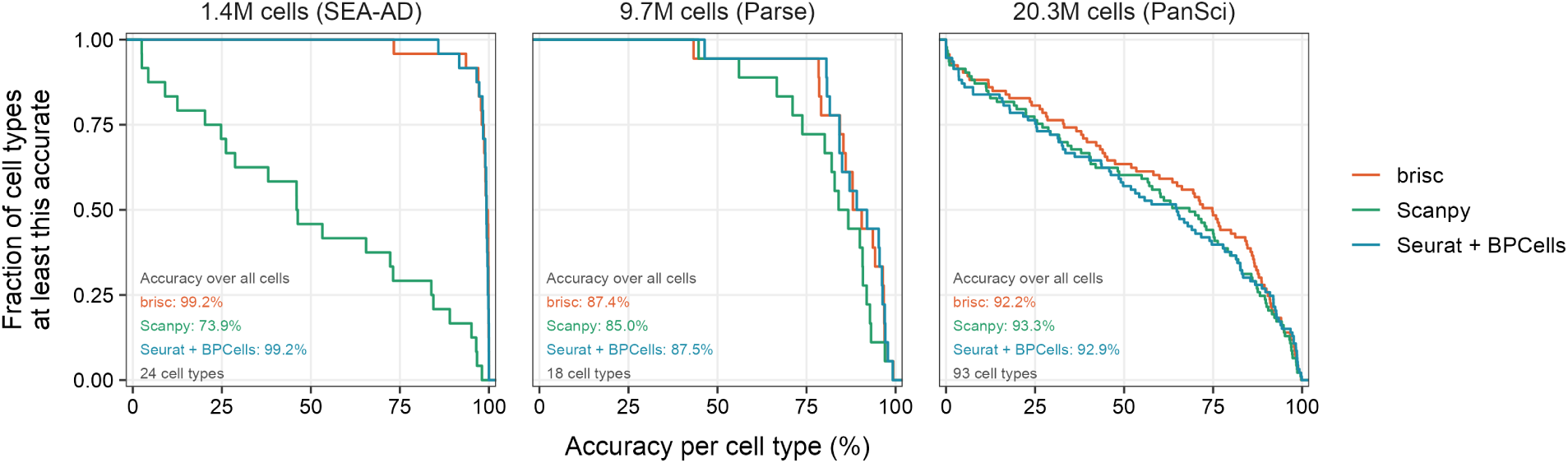
Accuracy of transferred cell-type labels. Fraction of cell types whose label-transfer accuracy meets or exceeds each value, per toolkit and dataset. A cell type’s accuracy is the percentage of its cells whose transferred label matches the author annotation, averaged over the three runs. Numbers in the bottom left of each panel show each toolkit’s overall accuracy across all cells and the number of cell types.

## Notes

https://github.com/briscverse/brisc

https://brisc.run

https://github.com/keon-arbabi/brisc-benchmarking

### Competing Interest Statement

The authors have declared no competing interest.

